# One-Pot Quantitation of Mycobacterial N-terminal Protein Acetylation Peptidoforms and Proteome

**DOI:** 10.1101/2025.03.30.646241

**Authors:** Daniel D. Hu, Simon D. Weaver, Bradley S. Jones, Patricia A. Champion, Matthew M. Champion

## Abstract

Isotopic labeling of proteins for quantitative proteomics is a popular technique to increase sample throughput and provide improved accuracy and precision for relative or absolute quantitation between samples. Derivatives of this technique are used to label protein and peptide N-termini for selective enrichment and analysis. We previously reported on a method to enrich and quantify protein N-terminal acetylation in model-system and pathogenic mycobacteria. Significant recent advancements in silica filter-based protein digestion have improved identification of proteins in bottom-up proteomics. However, these are not yet compatible with existing methods which detect and quantify protein termini and N-terminal modifications. Here, we present a one-pot method (OnePotNαTA) that incorporates silica filter digestion with protein N-terminal labeling and subsequent quantitation. This technique achieves high-density coverage of the N-terminome and obviates the need to enrich N-terminal peptides prior to analysis. OnePotNα TA identified 54% of the canonical proteome from whole-cell lysates of *Mycobacterium marinum* and eliminated biases in peptides identified by forgoing enrichment. This significantly reduced sample preparation time by ≥5-fold and preserves protein-level abundance measurements (LFQ) from the same injections and analysis. Analysis of a mutant strain of *M. marinum* lacking Emp1 (*Δemp1*)-an N-terminal Acetyltransferase required for efficient pathogenesis-identified 37 putative substrates of this enzyme. Additionally, analysis of the remaining peptides identified at least 34 proteins with alternate true N-termini, distinct from the canonical genome.

## Introduction

N-terminal acetylation (NTA) is a ubiquitous protein co- and post-translational modification in organisms. This modification is the covalent attachment of an acetyl group to the α-amine of a protein N-terminus and is distinct from ε-amino acetylation found on lysine residues. Analyses across humans, mice, and yeast show that 50-90% of identified eukaryotic proteins are N-terminally acetylated, including virtually all exported proteins.^1–3^ In humans, acetylation is mediated by six N-acetyltransferases (NATs) and is functionally associated with changes in protein stability/degradation, protein-protein interactions, and cellular translocation.^4,5^ In prokaryotes, NTA is less abundant and frequent.^6^ Previous studies have reported 10-30% of the proteome is N-terminal acetylated in *Escherichia coli and Acinetobacter baumannii*, respectively.^7–10^ Despite the relative infrequence of NTA in prokaryotes, bacteria encode significant acetylation machinery in their genomes. *Mycobacterium tuberculosis (M. tb)* for example encodes at least 47 acetyltransferases.^11^ N-terminal protein acetylation in bacteria has been shown to correlate with virulence, protein stability, signaling and physiological responses.^12^ The density of acetyl transferases in mycobacteria suggests a more focused role for their functions, in contrast to the ‘generalist’ strategy used by yeast and other eukaryotes^1-3,10,15^. Despite their prevalence and role in pathogenesis, N-terminal protein acetylation is understudied in part due to inadequate bioanalytical tools for census and characterization. ^13^

There are several existing approaches to quantitate protein NTA. One of the earliest techniques coupled with mass-spectrometry based proteomics to identify and quantitate NTA on a proteome-level was <u>Co</u>mbined <u>Fr</u>actional <u>Di</u>agonal <u>C</u>hromatography (COFRADIC).^14,15^ In this method, free protein N-termini are blocked with acetyl/deuterated-acetyl groups, followed by trypsin digestion. Proteins are fractionated by reverse-phase chromatography, followed by reaction with 2,4,6-trinitrobenzenesulfonic acid (TNBS) which blocks the remaining N-termini of ‘neopeptides’ produced from digestion. A second reverse-phase fractionation separates the hydrophobically distinct peptide groups, modulated by TNBS addition, resulting in enriched N-terminal peptide fractions that can be analyzed via LC-MS/MS^14,15^.

Several other techniques have been developed for N-terminal peptide analysis, virtually all incorporating methods of positive or negative enrichment.^16–19^ We previously developed and reported on a method that follows COFRADIC principles to quantify protein N-terminal acetylation from mycobacteria.^10^ Using filter-aided sample preparation (FASP) as the reaction chamber, protein N-termini were trideutero-acetylated, digested, and eluted for downstream enrichment by cation-exchange and nLC-MS/MS-MS analysis.^10^ This technique identified 45% of the canonical proteome in M. tb, and 26% of the N-termini were detected. However, these techniques are time-consuming, (5-days) onerous to perform on large sample sets, and error prone. The extended time-period of sample preparation introduces significant artifacts including pyro-glutamic acid formation, O-acylation, and excessive oxidation, which reduce sensitivity and required complex chemical and enzymatic steps to mitigate.^10,14,15^

We sought to develop an improved methodology that increases sample throughput without sacrificing N-terminal identification. We built and designed this method around incorporating immobilization of proteins using silica depth filters e.g. suspension trapping.^20^ Suspension trapping enables use of strong detergents (e.g. sodium dodecyl sulfate) prior to trypsin digestion, and has proven versatility in its ability to be modified and adapted to fit-for purpose approaches.^21–23^ Silica-depth filters are chemically compatible with most miscible and immiscible solvents facilitating derivatization of immobilized proteins, improving extraction, and contaminant-removal options. Silica-trapping requires lower sample preparation time compared to other SDS-compatible techniques such as filter-aided sample preparation (FASP), aligning sample throughput with iterative/temporal biological analysis of N-terminal acetylation. Our new method, coined OnePotNαTA, utilizes *in-situ* chemical-derivatization on suspension trapped (STrap) proteins, followed by digestion in the same chamber and forgoes enrichment of N-terminal peptides, prior to nLC-MS/MS. This approach virtually eliminates sample preparation artifacts and preserves quantitative Label-Free (LFQ) analysis of underlying changes in the responding proteome We apply this new technique to the analysis of the N-terminome of *M. marinum*. We analyzed the N-terminal proteome of an *M. marinum* mutant with a genetic deletion of *emp1*, a NAT with a recently defined role in virulence and known substrates.^35,36^ and compared it to WT *M. marinum*. We identified at least 34 novel protein termini, confirmed dozens of acetylated protein substrates of Emp1 and demonstrate that conservation of protein quantification distinguishes changes in abundance from changes in occupancy of the underlying PTM.

## Methods

### Materials

Ammonium bicarbonate (ABC), triethylammonium bicarbonate (TEAB), phosphoric acid, Sautons, 7H9, 7H11 media, Amicon Centrifugal Filters, and all other chemicals not listed were from Sigma-Aldrich (Burlington, MA). LC-MS-grade water, acetonitrile, acetone, and methanol were from JT Baker (Radnor, PA). Tris(2-carboxyethyl)phosphine (TECP), 150 mL Rapid-Flow Bottle Top Filters, and Micro BCA Protein Assay Kit was from Thermo Scientific (Waltham, MA). Iodoacetamide (IAM) and sodium dodecyl sulfate (SDS), and phenyl methyl sulfonyl fluoride (PMSF) were from VWR (Radnor, PA). Trypsin was from Promega (Madison, WI). S-Trap Minis were from Protifi (Fairport, NY). Hydrophilic-lipophilic balance (HLB) solid-phase extraction (SPE) cartridges were from Waters (Milford, MA). LC-MS/MS system used is a nanoElute 2 and timsTOF Pro 2 from Bruker Scientific (Billerica, MA). Tween 80 was from Fisher Scientific (Waltham, MA). Zirconia disruption beads were from Research Products International (Mount Prospect, IL).

### Synthesis of N-Hydroxysuccinimidyl esters (NHS) esters

The following synthesis was adapted from Staes et al. and Thompson et al.^10,15^ 1.77 g of N-hydroxysuccinimide (0.015 mmol) was mixed with 5.0 g of acetic anhydride-d_6_ (0.046 mmol) in a round-bottom flask with magnetic stir bar overnight (12-16 hours), with aluminum foil loosely wrapped around the mouth of the flask. The aluminum foil was then removed and reaction left to stir for an additional 12 hours, letting excess acetic anhydride evaporate. ∼10 mL increments of hexanes were added to the flask, stirred, and removed with rotary evaporation. Two additional washes of hexanes were performed, followed by two 10mL washes with anhydrous acetonitrile, and dried with rotary evaporation. No further purification was performed. 7.0 mL of anhydrous acetonitrile was added to the flask to dissolve NHS esters completely. 50 mg aliquots of NHS esters were pipetted into microcentrifuge tubes (141.9 µL per aliquot), followed by vacuum concentration until fully dry. Validation of reaction was tested by TLC (R_f_=0.2). and labeling of leu-enkephalin YGGFL.^15^ Aliquots were stored at -20°C until use.

### Growth of bacterial cultures

*M. marinum* cultures were grown in Sauton’s defined media to induce ESX-1 protein export into culture supernatants as described previously.^35,36^ Briefly, *M. marinum* colonies were picked from 7H11 plates, grown in 7H9 defined media (3 days) and subsequently diluted to an OD_600_ of 0.8 in 50 mL of Sauton’s media supplemented with 0.05% Tween 80. Cultures were grown at 30°C for 48 hours while shaking (200 rpm). Cells were grown for 48-hours and cell pellets and culture filtrates were harvested via centrifugation. Whole-cell lysates were subsequently generated by resuspending the bacterial cell pellet in 0.5 mL of sterile PBS plus protease inhibitor (PIC), 0.1mm zirconia disruption beads. Cells were then lysed using a Mini-Beadbeater (3×60 s on ice). Cell lysates were clarified by centrifugation at 12,000 x g for 12min. Lysates were transferred to new 1.5 mL microcentrifuge tubes and stored at -80°C until use. Culture filtrates were sterile filtered, supplemented with PMSF, and concentrated to ∼200 μL total volume using Amicon Ultra-Centrifugal Filters according to manufactures instructions with additions of PBS^36^. Protein concentrations of whole-cell lysates and culture filtrates were determined via the Micro BCA Protein Assay Kit prior to downstream experimental use. Samples were prepared in biological duplicate.

### Labeling and digestion of proteins

For a detailed protocol, see supplementary information. Briefly, 100µg of *M. marinum* lysate (<50µl) was precipitated in 1.3 mL of acetone (>7-fold addition) for 2 hours at -20°C. Samples were spun at 12,000 x g for 10 minutes to pellet the precipitant. Acetone was decanted to waste, and samples were vacuum-concentrated to dryness with no heat to remove residual acetone (∼10 min.).

Protein samples were resuspended in 10µL 100mM TEAB, 5µL 1M TEAB, 25µL 20% SDS. Samples were intermittently heated and vortexed to ensure all proteins were dissolved. 5µL of 100mM TCEP was added, followed by 10 minutes of incubation at 95°C. Samples were cooled, and alkylated with 5µL of 100mM IAM in darkness for 30 minutes. Samples were acidified with 5µL of 12% H_3_PO_4_, flocculated with 350µL 100mM TEAB in 90% methanol, and loaded on S-Trap Mini filters. Samples were spun at 1,200g for 30 seconds to pass liquid completely through filter columns. Columns were spun down with one wash of 150µL 100mM TEAB in 90% methanol, and one wash of 150µL 1:1 MeOH/CHCl_3_.

Acetoxy-d_3_-NHS ester aliquots (50mg) were resuspended in 518µL of 10% methanol in acetonitrile (v/v) to 600mM. 75µl was applied to the silica-filter incubated 30 minutes at room temperature, spun down, and incubated with a second 75µL addition for 30 min. Two washes of 150µL 50mM ABC in 90% methanol were added and spun through, to buffer-exchange and quench any residual unreacted NHS. Spin filters were placed in new collection tubes prior to digestion.

2µg of MS-grade trypsin was added to each sample in 160µL of 50mM ABC. Samples were incubated overnight at 37°C (minimum 4 hours). After digestion, samples were spun down to collect peptides, followed by two elution washes of 80µL 0.1% formic acid in water and one wash of 80µL 0.1% formic acid in 50% acetonitrile. Samples were vacuum-concentrated for 20 minutes at 50°C to evaporate acetonitrile, in order to prepare for desalting by solid-phase extraction (SPE).

### Solid Phase Extraction (SPE)

Samples were desalted using HLB SPE cartridges on a vacuum manifold. Cartridges were wetted with two additions of 500µL acetonitrile and equilibrated with three additions of 250µL 0.1% formic acid in water. Samples were loaded on SPE, followed by two washes of 250µL 0.1% formic acid in water. Cartridges were removed from manifold, placed over microcentrifuge tubes, and eluted in three additions of 200µL 0.1% formic acid in 50% acetonitrile. Samples were dried down and resuspended to a fixed concentration (500ng/µL) for LC-MS analysis.

### nLC-MS-MS/MS

Samples were analyzed using a Bruker nanoElute and timsTOF Pro LC-MS system with CaptiveSpray source. 90 min gradients were used on an Ionoptiks 25cm x 75µm, 1.6µm C18 column. For Emp1-related samples, a Pepsep XTREME C_18_ column (25cm x 150µm, 1.5µm) column was used with 120 min gradients (5% to 35% 0.1% formic acid in acetonitrile). PASEF-DDA acquisition method was performed using manufacturer’s default settings with the following exceptions: quadrupole low mass cutoff set to 80m/z, scan mode from 80-1700 m/z, and pre-pulse storage set to 8ms. Each biological replicate was injected in technical triplicate.

### Protein ID/Inference, Data Analysis

Raw (.d) files were searched using the PEAKS Online search engine (build 1.4.2020-10-21_171258). PEAKS DB analysis was performed using a custom modified protein database. The *Mycobacterium marinum* individual gene list (release 4; 5,453 entries) was downloaded from Mycobrowser (https://mycobrowser.epfl.ch/) and translated to amino acid sequences in R (v4.3.2) using seqinR (v4.2-30). All initial amino acids not translated as methionine (M) were manually changed to M (see Supplemental Figures 2 and 3). This proteome was saved as a .fasta file and used in the database search. Search settings included a semi-specific trypsin search with up to 3 missed cleavages which compensates for the loss of cleavage after lysine due to isotopolog-acetylation. A fixed modification of cysteine carbamidomethylation was added, along with the following variable modifications: acetylation (both light +42.01 and heavy +45.03) of peptide N-termini and lysines, deamidation of asparagine and glutamine, pyroglutamic acid formation from N-terminal glutamine and glutamic acid, and oxidation of methionine. Up to five variable modifications per peptide were allowed. All other settings and false discovery rate (FDR) control were set to manufacturer’s defaults.

### Peptide Analysis

Peptide lists were exported from PEAKS and analysis was performed in R (v4.3.2). For the genomic analysis, the same database from the PEAKS search above was used and an *in silico* tryptic digest was performed using the package cleaveR (v1.38.0). N-terminal peptides where the second amino acid was G, A, S, T, P, C, or V, had the initiator methionine removed.^24^ For the comparison with enrichment data, the supplemental information from Thompson et al.^10^ was downloaded as the worksheet entitled “Raw NTA Peptides” and the peptides identified in the tryptic digestion injections were analyzed. Isoelectric points were calculated with the seqinR (v4.2-30) package, ignoring N-terminal acetylation so that the identified peptide attributes could be compared directly with the genomic sequences. For all datasets, except to determine novel N-termini, peptides were considered N-terminal if they were observed at position 1 or 2 of the annotated protein sequence in the genome. Peptides of length 5-50 were considered. All plots were created with ggplot2 (v3.5.1).

### N-Acetyl Transferase Emp1 proteo-genetic analysis

Protein Label-Free Quantification (LFQ) and peptide database search .csv files were downloaded from PEAKS. All analysis was performed in R (v4.3.2). Differential protein expression analysis was performed with the packages QFeatures (v1.10.0) and limma (v3.56.2). Protein quantities were log transformed and normalized with the ‘center.median’ function in QFeatures. Technical and biological replicates (n=6) were all included, and a linear model was fit to the protein expression data using lmFit(), and empirical Bayes statistics were calculated with eBayes().^25^ This includes a Benjamini-Hochburg correction to produce adjusted p-values. 95% confidence intervals were calculated from these adjusted p-values.^26^ Proteins with adjusted p-values of less than 0.05 and log_2_ fold changes of > ±1.0 were highlighted as significantly changing. It is important to note that this significance combines biological and technical (analytical) variance, so is less biologically relevant than biological variance alone, but demonstrates the analytical precision of our methodology is sufficient to generate statistically significant results. Average normalized expression was plotted for each protein in WT vs each strain to generate differential expression plots.

Identified N-terminally acetylated peptides were filtered from the peptide data file and identified as heavy (+45.03) or light (+42.01). For each protein that had an identified light N-terminal acetylated peptide, the best flier peptidoform was identified by taking the peptidoform with the largest average area among all injections. The cognate heavy version of this peptidoform was identified for each protein. For each protein in each injection, a percent acetylation value was calculated by taking [light peptide area / (light peptide area + heavy peptide area)] ^*^ 100 (%). Peptides missing light or heavy peptides in any given injection were assigned 0% or 100% acetylated respectively. For each protein, an average percent acetylation and standard deviation was calculated for each strain by combining each biological and technical replicate (n = 6). Proteins that contained less than 3 non-NA percent acetylation values between the six injections of WT and Δ*emp1*/comp were discarded as too sparse. T-tests were performed between the WT and Δ*emp1* percent acetylation values. In cases where there was 0% acetylation in the Δ*emp1* strain and non-0% acetylation in the WT strain, the p-value was set as 0.001 (arbitrarily significant) for filtering purposes. %RSDs were calculated by taking standard deviation / mean for each percent acetylation value, and for percent acetylation of either 0% or 100%, %RSD was calculated by taking as the standard deviation / mean of either the light (100%) or the heavy (0%) peptide area. Acetylation was considered changing in response to deletion of *emp1* (MMAR_1839) if the following conditions were met: p-value of WT vs Δ*emp1* of < 0.05, higher percent acetylation in WT and Δ*emp1*/comp than in Δ*emp1*, and a change of percent acetylation between WT and Δ*emp1* of at least 20 absolute percentage values (e.g. 60% vs 39% included, but 20% vs 5% not included). The percent acetylation values for these proteins were plotted in each strain, with the size and shape of the point based on the %RSD value of the measurement. These proteins were highlighted in green on the protein differential expression plots.

Potential non-canonical (or misannotated) protein N-termini were identified by filtering out light N-terminally acetylated peptides at position 3 or later of the protein sequence. The same percent acetylation workflow described above was carried out on these proteins. Nine examples showing potentially non-canonical acetylated N-termini were extracted and plotted in the same manner as described above, with error bars showing the standard deviation of the percent acetylation (n=6). The point size is correlated with the number of non-NA values, with the smallest points representing 3/6 valid measurements and the largest being 6/6. Three of these examples showed strong *emp1* correlation (probable substrates of Emp1), three show weak correlation with *emp1*, and three do not correlate.

## Results/Discussion

We sought to develop a new method to quantify protein N-terminal acetylation that improves the time and number of steps necessary to process proteome samples at scale. Fundamentally most bottom-up proteomics methods for N-terminal analysis rely on distinguishing N-termini from neo-termini generated during digestion. This is typically achieved by selective chemical labeling, digestion in-solution; and a combination selective enrichment/depletion/desalting columns are applied.^15^ We previously published work to identify and quantitate protein N-terminal acetylation utilizing filter-aided sample preparation (FASP) as the microreactor for preparation.^10^ Proteins are spun onto a molecular weight cutoff filter, enabling chemical additions like detergents and reducing agents to be removed after incubation. This enabled selective addition and removal of chemical and enzymatic labeling reagents such as NHS esters and pGAPase and Qcyclase enzymes. While FASP enabled selectivity in introducing and removal of deleterious reagents from the protein sample, it is time consuming with significant hands-on manipulation and centrifugation at every step. Batch-to-batch variability and robustness are poor-to-moderate, as different samples affect the speed in which filters are completely spun through and filter quality control is uncertain.^20,27^ This requires careful supervision of centrifugation steps to ensure each sample is similarly treated to prevent excessive drying or membrane failure

Here, using OnePotNαTA (outlined in Figure 1), we utilized commercial suspension trapping (S-Traps) with silica depth filters to achieve superior selectivity while reducing hand-on and centrifugation inputs to a fraction of the time (<30 sec per wash). Spin columns are composed of a silica matrix with significantly lower back pressures during centrifugation compared to cutoff filters used in FASP. Broadly, proteins solubilized by SDS are flocculated in a high ionic-strength methanolic solution, creating a suspension of precipitated proteins that associate with and are immobilized by the silica filter. SDS is removed by addition of buffered methanol leaving the immobilized protein amenable to proteolytic digestion. Past comparisons between the suspension trapping and FASP filter techniques demonstrated negligible differences in protein identification fidelity.^20-22,28^

**Figure 1.**
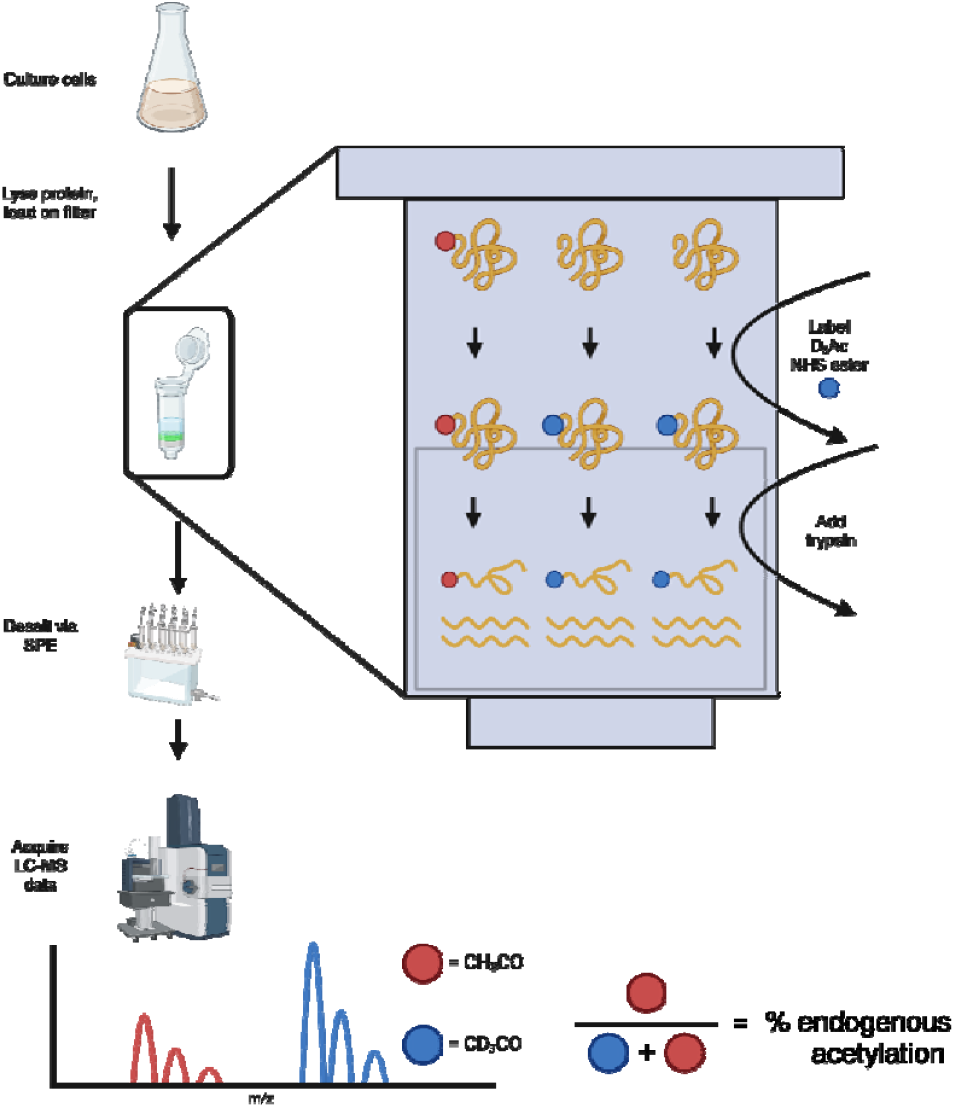
Workflow of one-pot method to quantify N-terminal acetylation. Extracted proteins are reduced, alkylated, and immobilized on silica depth filters. Spin columns act as a one-pot bioreactor, where trideutero acetoxy-NHS esters other labeling reagents, wash/quench reagents, and trypsin can all rapidly pass through for selective addition and removal of reagents as needed. Free protein N-termini (And typically K) are labeled with stable heavy isotope acetyl groups, enabling mass differentiation of endogenous acetylation at and termini the proteome/peptitoform level.

We hypothesized that the silica environment used for trapping and digestion would be well-suited to chemical label proteins prior to digestion. We designed this method to chemically label intact protein N-termini *on-filter* between wash steps. This enabled the same benefits and convenience of one-pot bioreactors to selectively introduce reagents and wash out unwanted contaminants, reducing sample preparation time from a 5-day process to less than 6 hours.

Labeling on-filter reduces undesirable side-reactions. One concern when using NHS esters to label protein N-termini is the frequency of O-acylation of serine, threonine, tyrosine amino acid side chains.^29,30^ N-terminal peptides with variably modified O-acyl groups reduces sensitivity, confounds the search space and makes endogenous NTA quantitation challenging. O-acylation is reversible by addition of hydroxylamine at high pH or boiling samples to thermally reverse the labile O-acyl bond.^31,32^ Both of these approaches introduce artifacts, oxidation and potential non-enzymatic cleavage at NG and other amino acids. Addition of alcohol during labeling will significantly reduce protein O-acylation, which is commonly done with peptide isobaric tagging, but is generally not compatible with FASP and protein-labeling due to solubility and filter breakdown. We hypothesized that our approach (Figure 1) using the suspension trap to derivatize after protein binding would eliminate N-terminal enrichment artifacts. Proteins trapped on silica filters are compatible with organic solvent additions (e.g. MeOH) which eliminates O-acyl formation and excess reagent are readily removed after labeling, which further reduces chemical artifacts. The 40-fold reduction in preparation time also significantly reduces the formation of base catalyzed and time dependent artifacts like pyro-glutamate formation and oxidation (e.g. M, W, H) residues. Frequency of N-pyro glutamic acid formation from E,Q and oxidation of M were comparable in frequency to that observed from unlabeled/nominal tryptic digests of *M. marinum* whole-cell lysates (data not shown).

We also switch the wash buffer after labeling switches from a compatible 3° amine (TEAB) to a 1° amine) ammonium bicarbonate, quenching any residual unreacted NHS esters. This approach yielded the greatest proteome coverage while minimizing off-target labeling.

We tested our OnePotNαTA method using whole cell lysate samples derived from cultures of *Mycobacterium marinum*. *M. marinum* is a model organism used to study *M. tuberculosis*, as it possesses high conservation of virulence pathways while having higher growth rate and lower risk to human health. The number of proteins and peptides identified in OnePotNαTA are summarized in Table 1. From a total canonical annotated proteome of 5,481 proteins (Mycobroswer release 4), 3,644 proteins were identified at a <1% FDR. These proteins were inferred from 65,395 peptide identifications, of which 2,536 were N-terminal (P1’/P2’). 2,464 peptides were N-terminally acetylated, of which 756 peptides were detected with isotopically light acetylation (endogenous acetylation) while 3,547 peptides were identified with d3-heavy label. The coverage of the proteome identified is shown as Figure 2. 53.7% of the proteome was identified while 24% of all N-termini from the proteome were identified. This is in good agreement with previous studies showing 10-30% of prokaryotic proteins being N-terminally acetylated.^7–10^

**Table 1.** Identification of proteins and peptides in OnePotαNTA.

|  |  |
| --- | --- |
| Proteins in proteome | 5,481 |
| Proteins identified | 3,644 |
| Peptides identified | 65,395 |
| N-terminal peptides identified (P1'/P2') | 2,536 |
| Proteins w/ N-termini identified (P1'/P2') | 951 |
| N-terminally acetylated proteins | 2,464 |
| N-terminally acetylated peptides | 4,363 |
| N-terminally light acetylated peptides | 756 |
| N-terminally heavy acetylated peptides | 3,547 |

**Figure 2.**
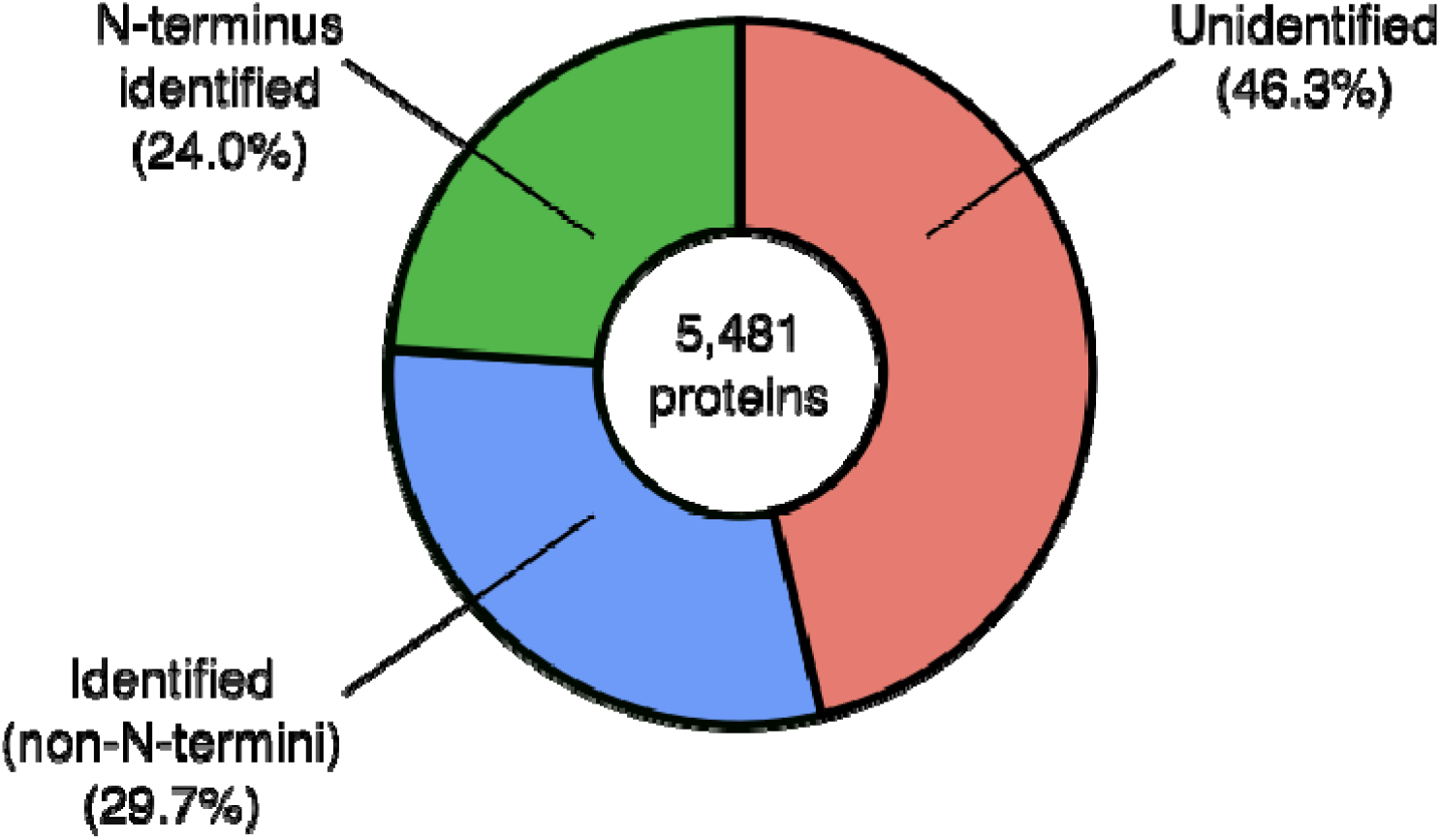
Coverage of proteins and N-termini identified. *M. marinum* annotated proteome contains 5,481 protein entries (Mycobrowser). 53.7% of the canonical proteome was identified, while 44.7% of the identified proteome contained N-terminal peptides.

Curation of N-terminal amino acids in database improves N-terminal peptide identifications in mycobacteria One discrepancy we discovered in database-based search was the presence of mistranslated protein databases in curated genome browsers at the P1’ position. Several genome browsers exist for mirobes, pathogens and other genera. For mycobacteria, Mycobrowser is a platform where gene and protein databases are annotated and updated with the most recent literature.^33^ Mycobacteria, encode many proteins with alternative start codons (non AUG). To scientific understanding these are not generally translated. Significant numbers of protein entries as translated in Mycobrowser annotate the non-canonical start codons (GUG->Val, UUG->Leu) to the N-terminus rather than substituting them with initiator formyl-methionines, comprising 29% of the proteins (1,595/5,481). Uniprot (https://www.uniprot.org/) by contrast to the best of our knowledge does not do this.

In order to test the impact of non-canonical translation initiation, we use a custom .fasta file in which N-terminal valines, leucines are were correctly substituted with methionine. Comparative database searches against the LC-MS raw data show a modest but significant improvement in the number of PSMs when searching using a database with all N-terminal amino acids replaced with methionine (SI Figure 2,3). The relative amount of detected N-terminal peptides for proteins (32%) starting with GUG is closer to the overall genome (29%) than those detected without using the custom database (17%) indicating that the detected N-terminome with this modified database better matches the expected N-terminome. (SI Figure 3). While minimally important for general bottom-up proteomics, this type of careful database curation and review is important for N-terminomics studies where a high percentage of N-terminal peptides are impacted, and should be noted by researchers studying non-canonical translation, translation initiation, and N-termini.

OnePot*α*NTA is not Isoelectrically (pI) Biased

**Figure 3.**
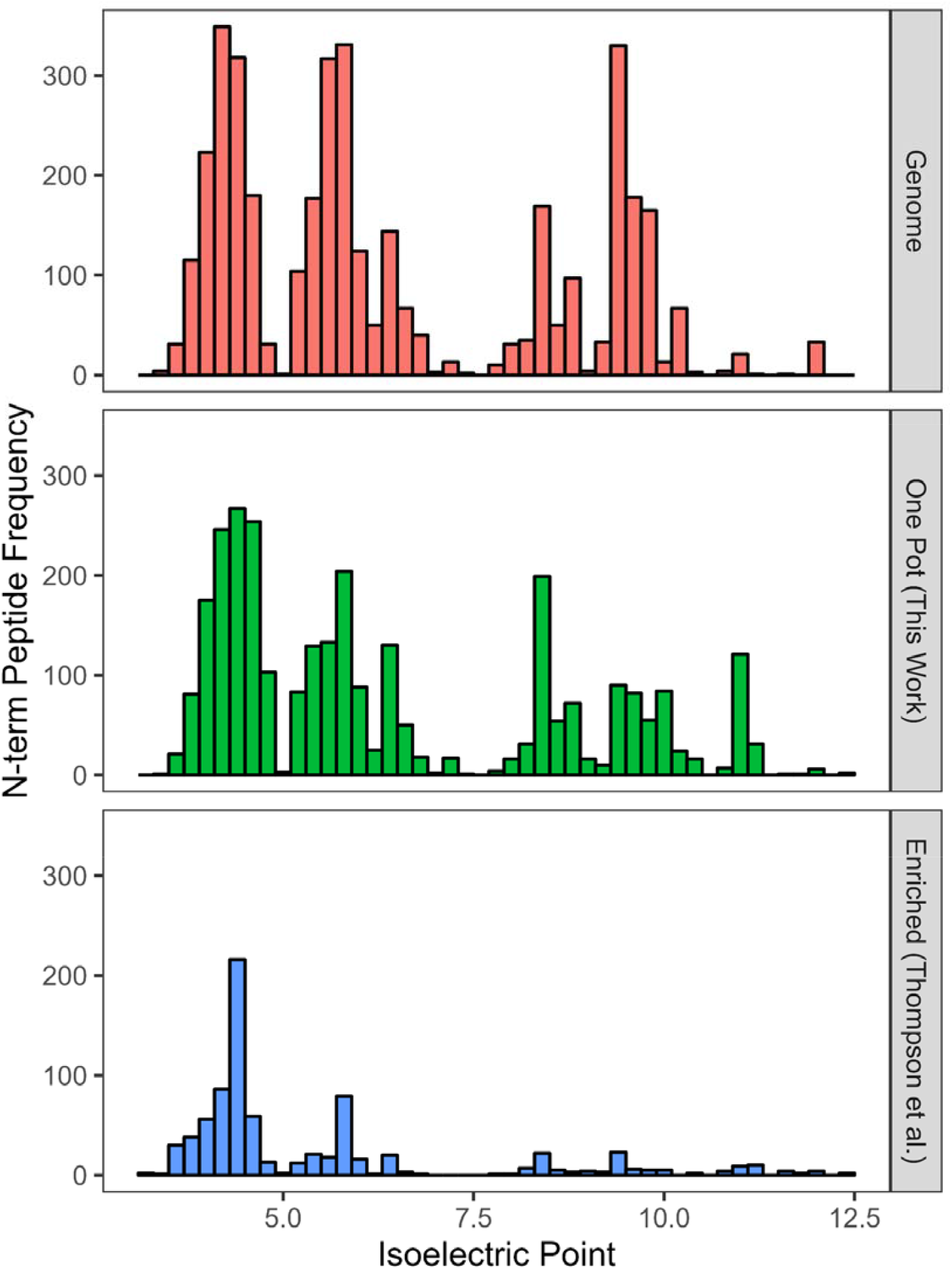
Histogram of isoelectric points of the identified N-terminal peptides. Top panel contains N-terminal peptides from an *in silico* tryptic digest of the *M. marinum* genome (red). Middle panel contains N-terminal peptides identified in our OnePotαNTA method (green). Bottom panel contains N-terminal peptides identified in a cation-exchange enrichment-based method as described by Thompson et al. (blue).

We hypothesized that the OnePotNαTA would avoid biases that occur in N-terminal enrichment methods. Staes et al. and our previous data observed that histidine-containing protein N-termini were poorly recovered from cation-exchange N-terminal enrichment due to the positive charge on the histidine (pKa = 6.0) mimicking the positively charged amine of neopeptides from trypsin. To test this, we compared isoelectric point distributions of our identified peptides to an *in silico* tryptic digest of the *M. marinum* reference proteome as well as our previously published cation-exchange NTA method from Thompson et al.^10^ Figure 3 shows the distribution of N-terminal peptide isoelectric points between the canonical genome, our OnePotNαTA approach, approach previously described by Thompson et al.^10^ When comparing against the enriched approach, our OnePotNαTA technique has an N-terminal peptide isoelectric point distribution matching closer to the canonical N-terminome than to the enriched sample. Enriched peptides are more acidic, as basic N-terminal peptides are likely to bind to the strong cation exchange resin. This leads to lower recovery of total N-terminal peptides, despite enriching the relative amount of N-terminal peptides. While enrichment is typically performed to increase sensitivity of identifying peptides of interest, we found no significant penalty in N-terminal identifications when foregoing ion exchange chromatography enrichment. Improvements to data density and proteome coverage granted by newer mass spectrometers enables this enrichment-free method that was previously infeasible.

Differences in isoelectric point distribution between our OnePotNαTA method and the canonical genome led us to investigate the specific amino acid distribution between the two. Figure 4 shows an amino acid histogram comparing between OnePotNαTA and the reference genome. Peptides from the OnePotNαTA method have similar amino acid distributions to the reference genome. Previous approaches to quantifying NTA required use of enzymatic treatments to eliminate formation of N-terminal pyroglutamic acid, and fully oxidaze Methionine ridues, which effectively serve as false positives from the enrichment process.^10^ Our OnePotNαTA approach foregoes enrichment, and preserves distribution of glutamic acid residues to the genomic reference. OnePotNαTA yielded more arginine-containing neopeptides and lysine-containing N-terminal peptides relative to the genome. This is due to the OnePotNαTA method labeling lysine residues with acetyl groups, lengthening the N-terminal peptides generated by trypsin which predominantly cleave C-terminal to arginine. This is consistent with enrichment methods being bias against basic N-terminal peptides due to their high pK cation nature being excluded from cation-exchange enrichment.

**Figure 4.**
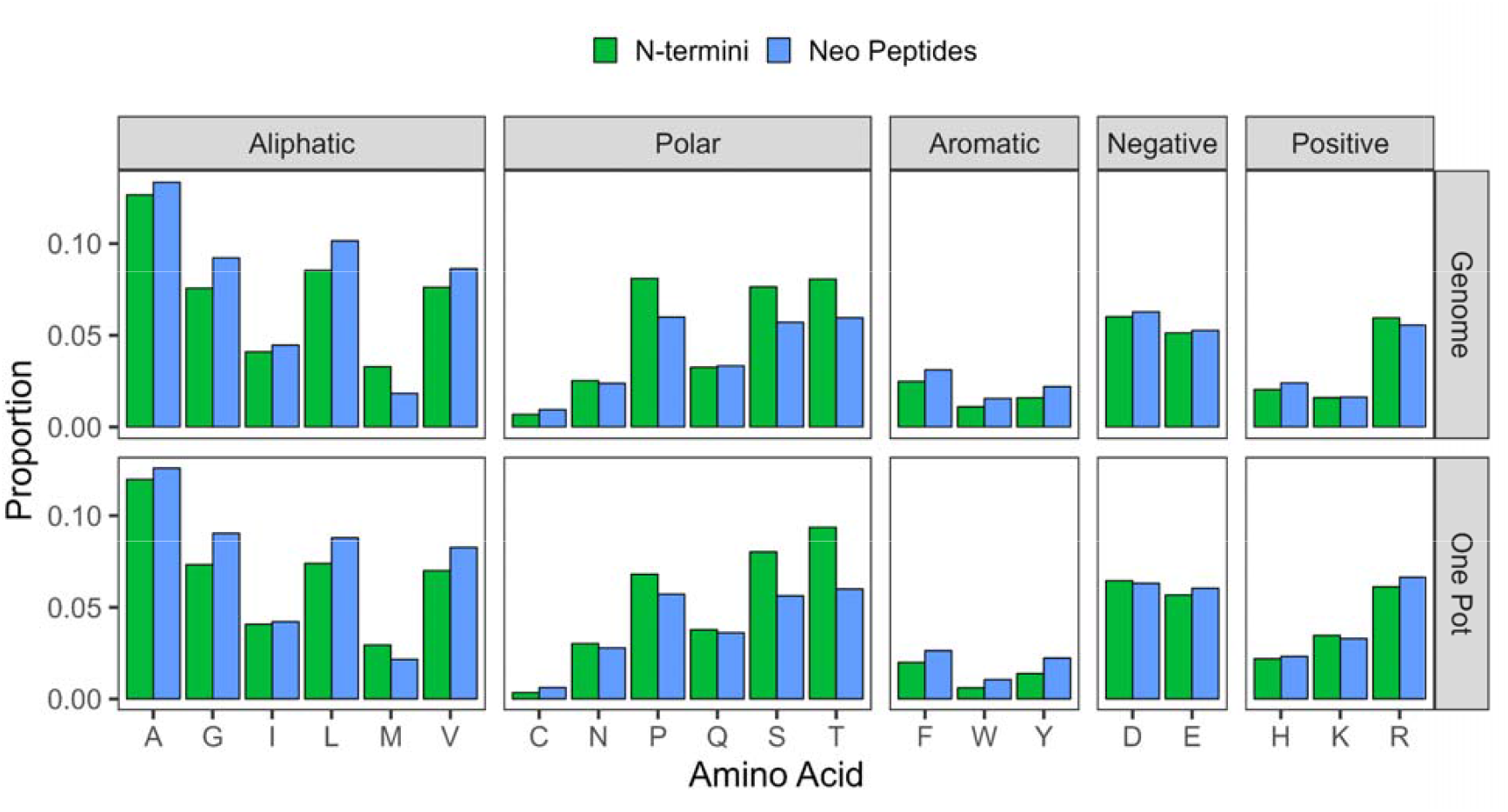
Amino acid distribution of peptides between OnePotαNTA and the canonical genome. Amino acids are separated by R-group category (columns). Green bars denote N-termini peptides while blue bars denote non-N-terminal neopeptides.

Both genome and OnePotNαTA show similarities in aliphatic and polar amino acid distributions. Polar amino acids are more abundant in N-terminal peptides than in neopeptides, while aliphatic amino acids are more abundant in neopeptides, with methionine being the exception. This is due to the ubiquity of methionine initiating all te protein translation processes, and its imperfect excision by methionine amino peptidease (MAP). Polar and aliphatic amino acids are considered to be stabilizing by the N-End Rule.^24,34^ Curiously, several stabilizing amino acids (Ala, Gly, Met, Val) were more prevalent in the neopeptides rather than the N-termini. This could be due to the amino acids’ contribution to structure in the middle of the sequence, and not the amino acids contributing less to the degradation rules of N-termini.

Our findings on the amino acid compositional differences between peptide N-termini and neopeptides demonstrates that the two groups form unique populations within the peptidome. Thus, we sought to further characterize chemical differences between the two. Supplemental Figure 4 shows the isoelectric point distribution between the peptide species. Top panels show all peptides, middle panels show N-terminal peptides, and bottom panels show neopeptides. Left panels are peptides generated from an *in silico* digest of *M. marinum* proteins, whereas right panels are peptides generated from our OnePotNαTA technique. These data show that while acidic N-termini and neopeptides are very similar in distribution, there are fewer basic neopeptides relative to N-terminal peptides. Isoelectric point distribution of peptides in our OnePotNαTA technique are similar to the genome-derived reference, albeit at a reduced dynamic range.

Similar comparisons were made using peptide mass and length distributions, shown in Supplemental Figures 5 and 6. No significant differences in mass or length were found between N-terminal peptides and neopeptides. Our OnePotNαTA technique did not recover all of the lower mass/length peptides found in the genome. This is a common occurrence in bottom-up proteomics, due to the optimized range of detection in the mass spectrometer and known biases against small peptides in all filter-based sample preparation approaches.^10,20-23^

**Figure 5.**
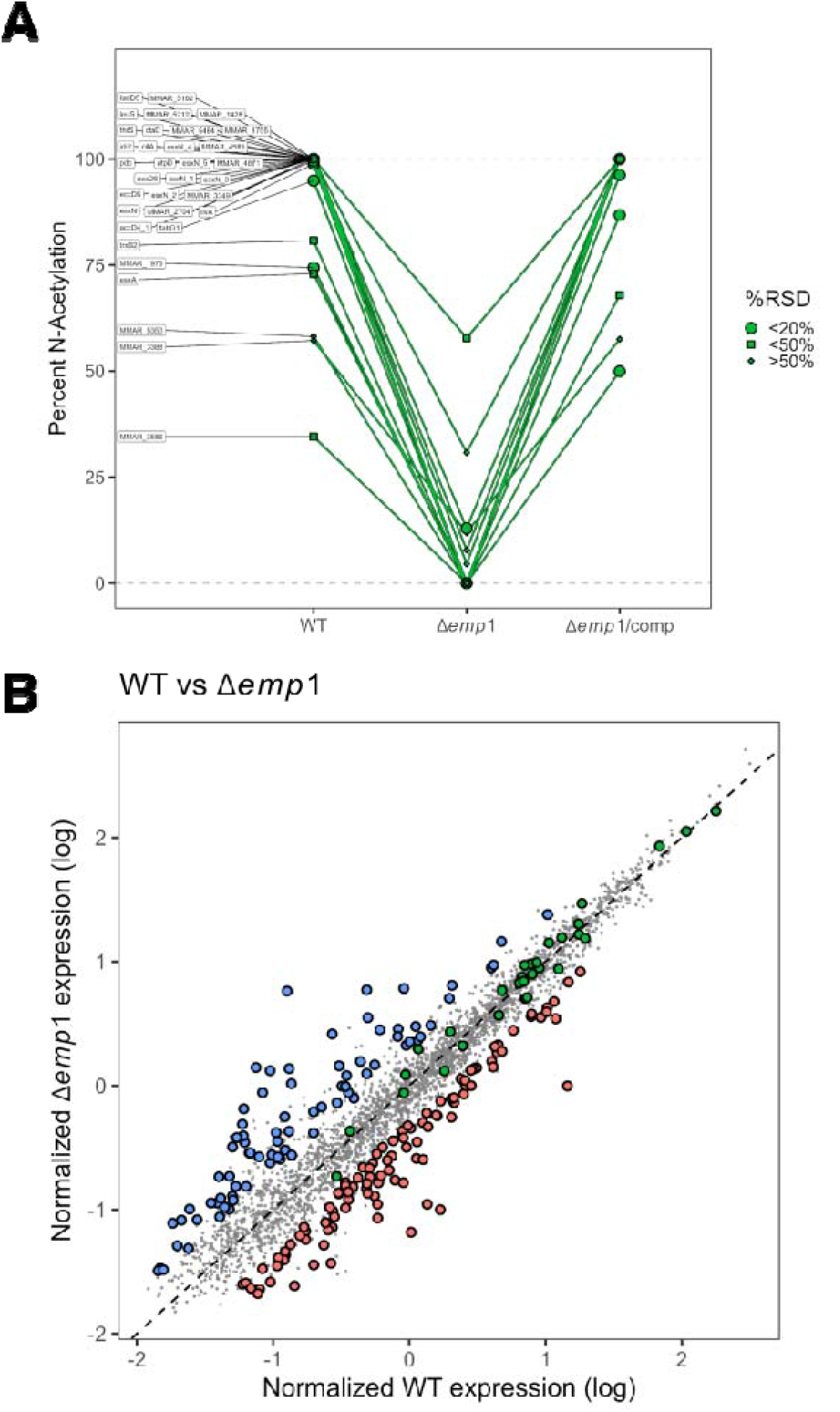
(A)N-terminal acetylation of proteins across three samples: wild type, Δ*emp1*, Δ*emp1* with complement. Point size is scaled to relative standard deviation (see legend). Data was filtered to only show proteins that exhibit a loss or attenuation in the deletion, followed by restoration in the complement sample. (B)Log-log plot of protein abundance between wild type and Δ*emp1* samples. Points in green denote proteins with differential N-terminal acetylation across samples. Blue and red points denote proteins with a fold-change greater than ±1, and a B-H adjusted p-value <0.05.

**Figure 6.**
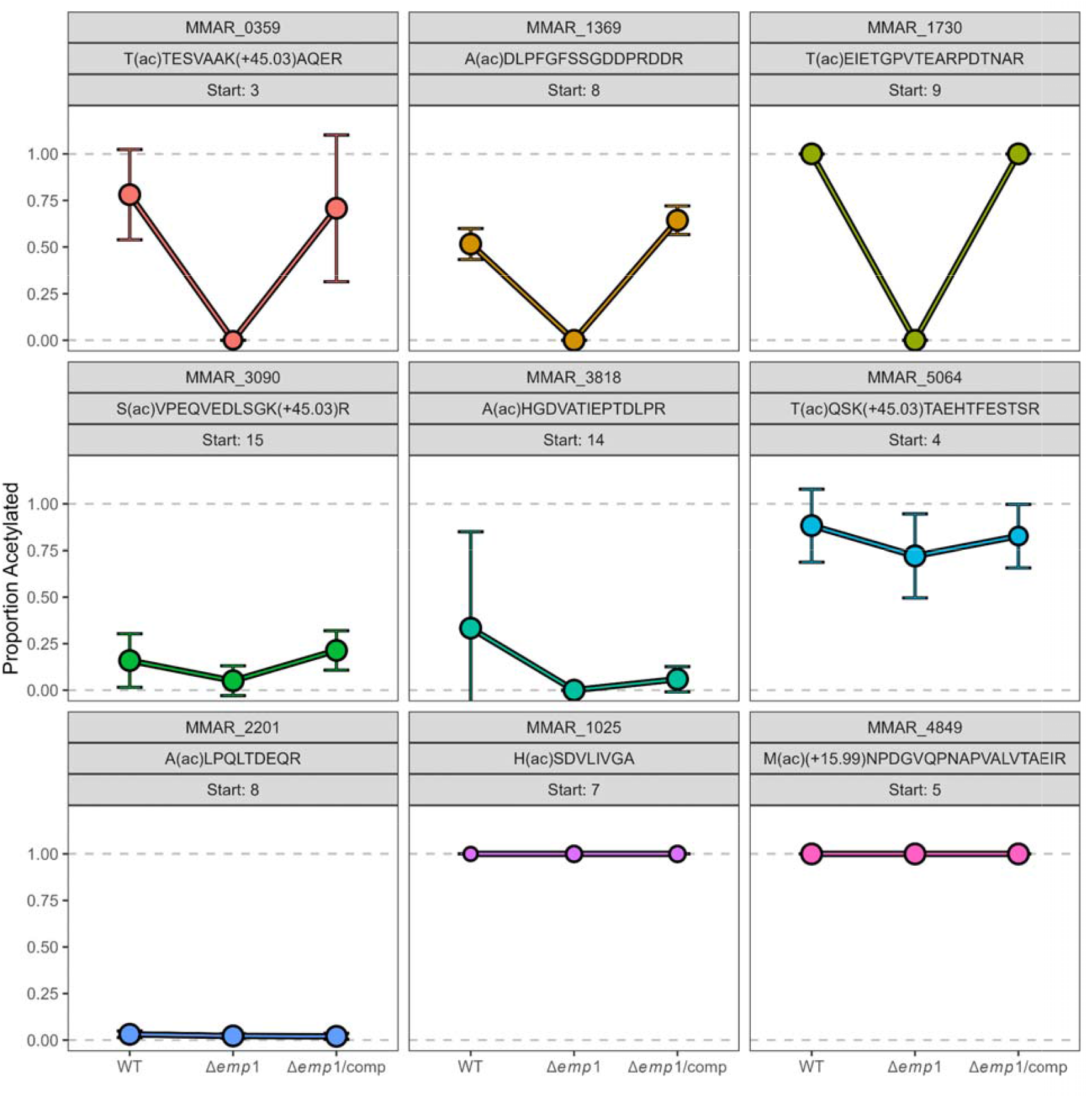
N-terminal acetylation of non-canonical start sites of proteins exhibiting strong (top three), weak (middle three), and no acetylation response (bottom three) to protein Emp1. Proteins with high Emp1-dependence show elimination of endogenous acetylation in the Δ*emp1* sample, and restoration in the complement.

We next compared the differences in LC-MS retention time between N-terminal peptides and neopeptides (Supplemental Figure 7). N-terminal peptides are scarcer and show greater stochasticity in identification across the gradient (panel B). A broad elution band of neopeptides appear between 70-80 minutes. N-termini exhibit higher density later in the gradient, whereas neopeptides distributed more evenly across the range of the gradient. Overall, neopeptides elute earlier in the gradient than N-terminal peptides. We hypothesize that this population is peptides for which labeling (due to blocked K/missed cleavage, and decreased polarity increases their relative detectability compared to canonical proteotypic peptides which mimic the distribution observed from 0-70 minutes.

### OnePotNαTA allows for simultaneous N-terminal measurement and protein quantification in an acetyltransferase mutant

We sought to apply our OnePotNαTA method on a set of samples with high biological relevance. We previously reported on identifying protein Emp1 as the acetyltransferase responsible for N-terminal acetylation of EsxA, a protein essential for virulence in *M. marinum*, and conserved in M. tuberculosis, the causative agent of human tuberculosis. EsxA is a substrate of ESX-1, a conserved export machinery in mycobacteria and acetylation of EsxA promotes virulence of M. tuberculosis in vitro.^35,36^

Additionally, we identified 22 putative substrates of Emp1 using bottom-up proteomics with label-free quantitation (LFQ).^35,36^ These data however, were not directly quantitative and relative abundance of acetylation could not be measured. We applied OnePotNαTA to these samples to demonstrate improvements not only in the detection and relative abundance of acetylation of the N-terminal acetylome, but also simultaneous quantification of the general proteome.

Figure 5 demonstrates how changes in protein N-terminal acetylation may be masked when only performing protein-level LFQ. Data in panel A are filtered to include only proteins in which N-terminal acetylation is attenuated or eliminated in the deletion strain, but restored in the complement. Y-axis represents the percentage of endogenous acetylation for each protein, calculated by light / (light + heavy). Panel B shows protein abundance between wild type and deletion samples; blue and red points denote proteins with a statistically significant fold-change greater than log_2_±1. Proteins that show differential N-terminal acetylation (green) do not show differential protein abundance. These data demonstrate that N-terminal acetylation is discontinuous with changes in protein abundance, and emphasizes the importance of analyzing both changes in NTA as well as protein quantitation. Using this method led us to identify 34 proteins that are Emp1-dependent for N-terminal acetylation (absent in mutant and restored in complement) and thus substrates of the enzyme (Supplemental Table 1), an increase of 55% over our label-free analysis. Six of these proteins are gene duplication paralogs (EsxN, EsxN_1 – EsxN_5) with the same N-terminal peptide sequence, meaning that there were 29 unique N-terminal amino acid sequences with Emp1-dependent acetylation.

### OnePotNαTA identifies putative non-canonical protein N-termini

As OnePotNαTA distinguishes N-terminal peptides with both light (+42 Da) and heavy (+45 Da) acetylation prior to tryptic digestion we surmised it would also identify novel N-termini similar to other N-terminally enriched studies, and complementary to 6-frame translation analysis. We analyzed our data to identify and reannotate true protein N-termini based on which amino acid position the acetylation (both heavy and light) occurs on. Figure 6 shows examples of proteins with N-termini distinct from the genome annotated translational start. Headers denote the position of the measured N-terminus in this dataset relative to the canonical N-terminus in the theoretical proteome. For example, the detected N-terminus of MMAR_1730 (B2HII5) (Figure 6, top right) is found at position 9 of the protein sequence in the database. The genome encoded N-terminus VIGNDRWV<u>T</u>EIETGPVTEA… Where we identify T9 as acetylated. This is not a signal sequence or similar cleavage event. The likely correct N-terminus is V(fMet)TEIETGPVTEA… We hypothesize that for this particular case translation initiated at canonical position 8, followed by initiator methionine cleavage and acetylation. The top three panels contain proteins that demonstrate a strong pattern of Emp1-dependent acetylation at these non-canonical N-termini; endogenous N-terminal acetylation is eliminated in deletion and restored in complementation. Abundant identification of both heavy and light N-terminally acetylated peptides not only aids in determining Emp1-dependent acetylation, but also confidence in true peptide N-termini. This is because both light and heavy N-terminal acetylation was identified in the same amino acid across multiple samples. Panels in the middle of Figure 6 demonstrate weaker Emp1-dependent acetylation. Endogenous N-terminal acetylation is attenuated in the deletion sample, but not outright eliminated. Panels in the bottom demonstrate proteins with noncanonical N-termini without Emp1-dependence. Our identification of noncanonical N-termini is not exclusive to Emp1-dependence, and is applicable across the entire proteome. In total, we identified at least 34 proteins with putative N-termini different from the genomic canon (Supplemental Table 2). Three of these showed strong Emp1 dependent acetylation, bringing the total number of protein products with Emp1 dependent acetylation (canonical and non-canonical) to 37, which have 33 unique N-terminal peptides.

## Conclusions

In summary, we developed a method of quantifying protein N-terminal acetylation that preserves protein-level abundance by foregoing peptide enrichment and improves sample throughput. We identified 53.7% of the canonical proteome in *Mycobacterium marinum*, of which 44.7% proteins contained N-terminal peptides. Our method recovers a higher distribution of basic N-terminal peptides than previously reported, and falls closer in line with the proteome. We identified biases in amino acid distribution between N-terminal peptides and neopeptides, prevalent with both OnePotNαTA and the reference proteome further demonstrating the decrease in bias with OnePotNαTA over enrichment methodology. We applied our method to samples of biological interest using Δ*emp1* and complement strains and demonstrated changes to N-terminal acetylation are not correlated with changes in protein abundance. We identified 34 proteins that show strong Emp1 dependent N-terminal acetylation. OnePotNαTA allows for the simultaneous measurement of both protein level LFQ and N-terminal acetylation, decreasing the number of required MS injections by half if both metrics are of interest to an experiment. We used our N-terminal acetylation data to map and reannotate true protein N-termini, identifying 34 proteins with putative noncanonical termini.

## Acknowledgements

The authors would like to acknowledge Dr. Boggess in the Notre Dame Mass Spectrometry and Proteomics Facility and the National Institutes of Health R01AI106872 (P. Champion) T32GM075762 (S. Weaver) for funding.

## Graphical Abstract

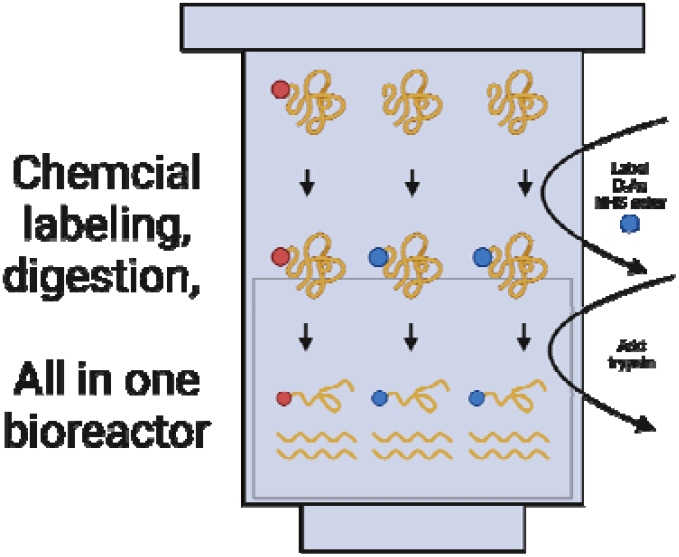

