## Supplementary material for "One-Pot Quantitation of Mycobacterial N-terminal Protein Acetylation Peptidoforms and Proteome": SI_Data

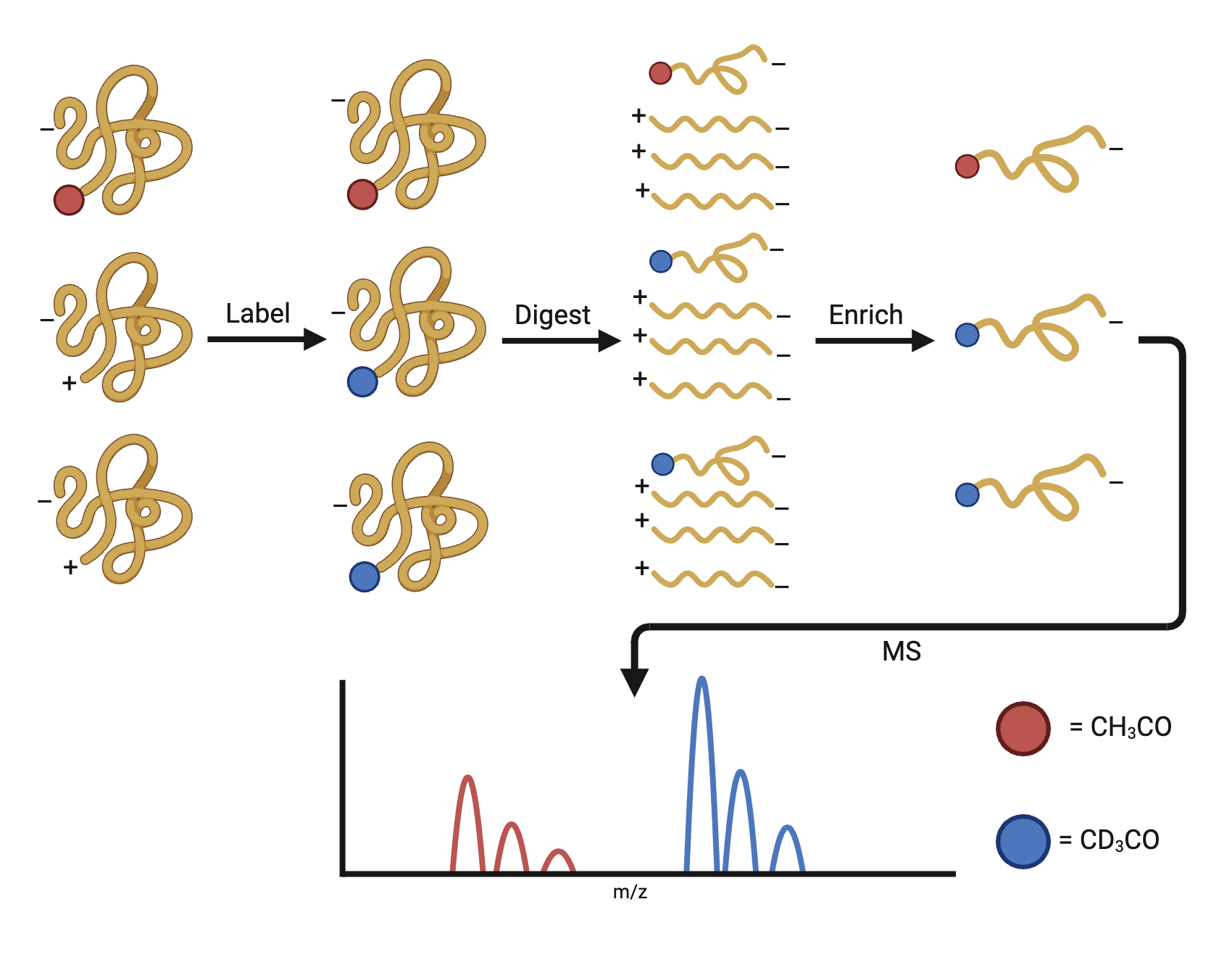

**Supplemental Figure 1.** Diagram of enrichment-based method to quantify N-terminal acetylation. Proteins are labeled with stable isotopically heavy acetyl groups, followed by digestion. Neopeptides (non-N-terminal) have a charge difference from N-terminal peptides, and are depleted by enrichment via strong cation exchange chromatography. Resulting N-terminal peptides are measured by LC-MS/MS analysis.

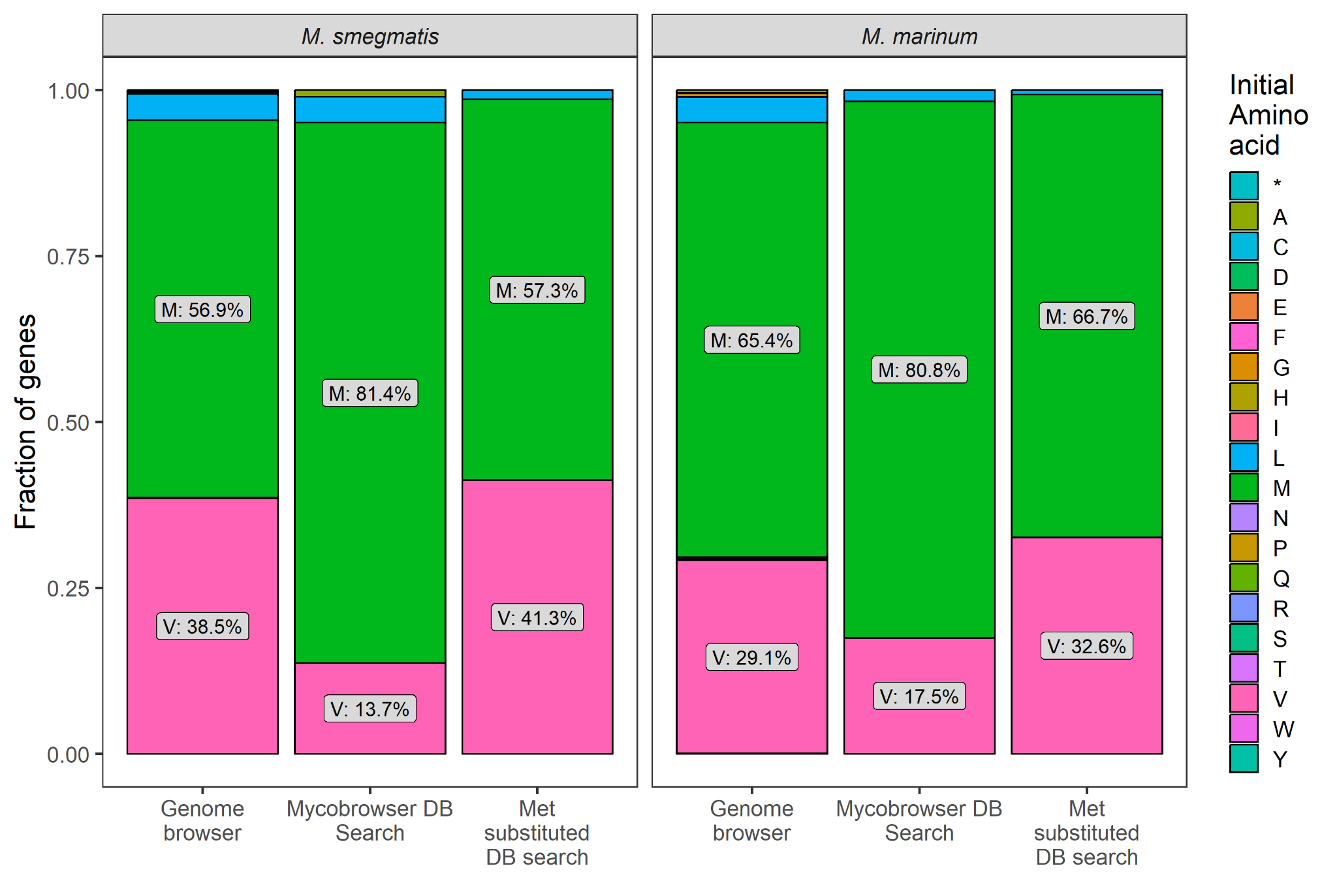

**Supplemental Figure 2**. Relative abundance of genes beginning with each amino acid. For each panel, the bar in the middle is protein N-termini identified in the raw database search strategy and the bar on the right is protein N-termini identified in the met-substituted database search strategy. These can be compared against the bar on the left representing the genome, which comes from Mycobrowser. The left panel is *M. smegmatis* and the right panel is *M. marinum*. Initial amino acid is based on the genome translated version. In the Met substituted database search all initial amino acids are searched as starting with Met

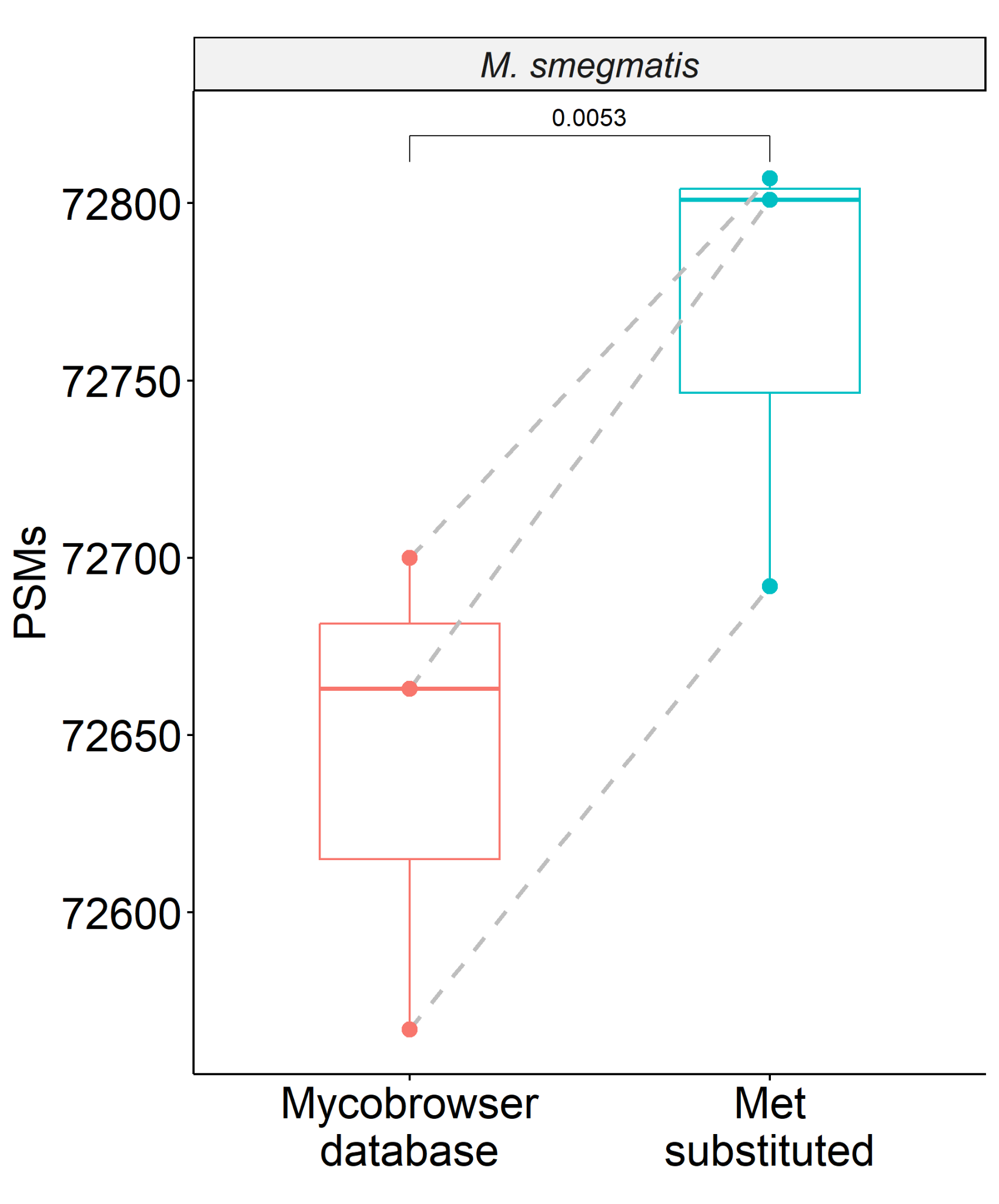

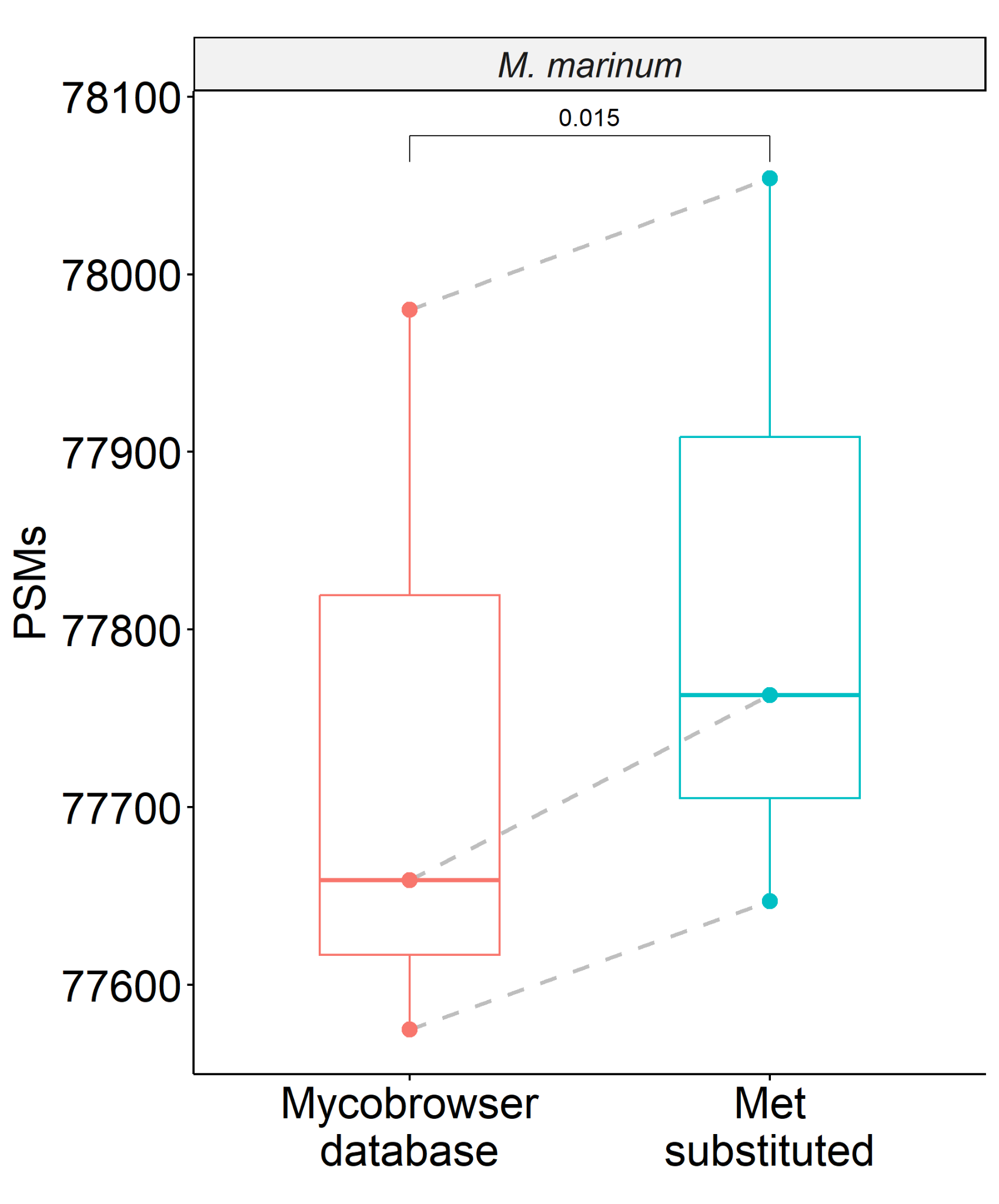

**Supplemental Figure 3**. Total PSM comparison from two database search strategies, one with N-terminal methionine substitutions (Met substituted) and one without (Mycobrowser database). P-values shown from paired t-test performed in R (n=3 for both species).

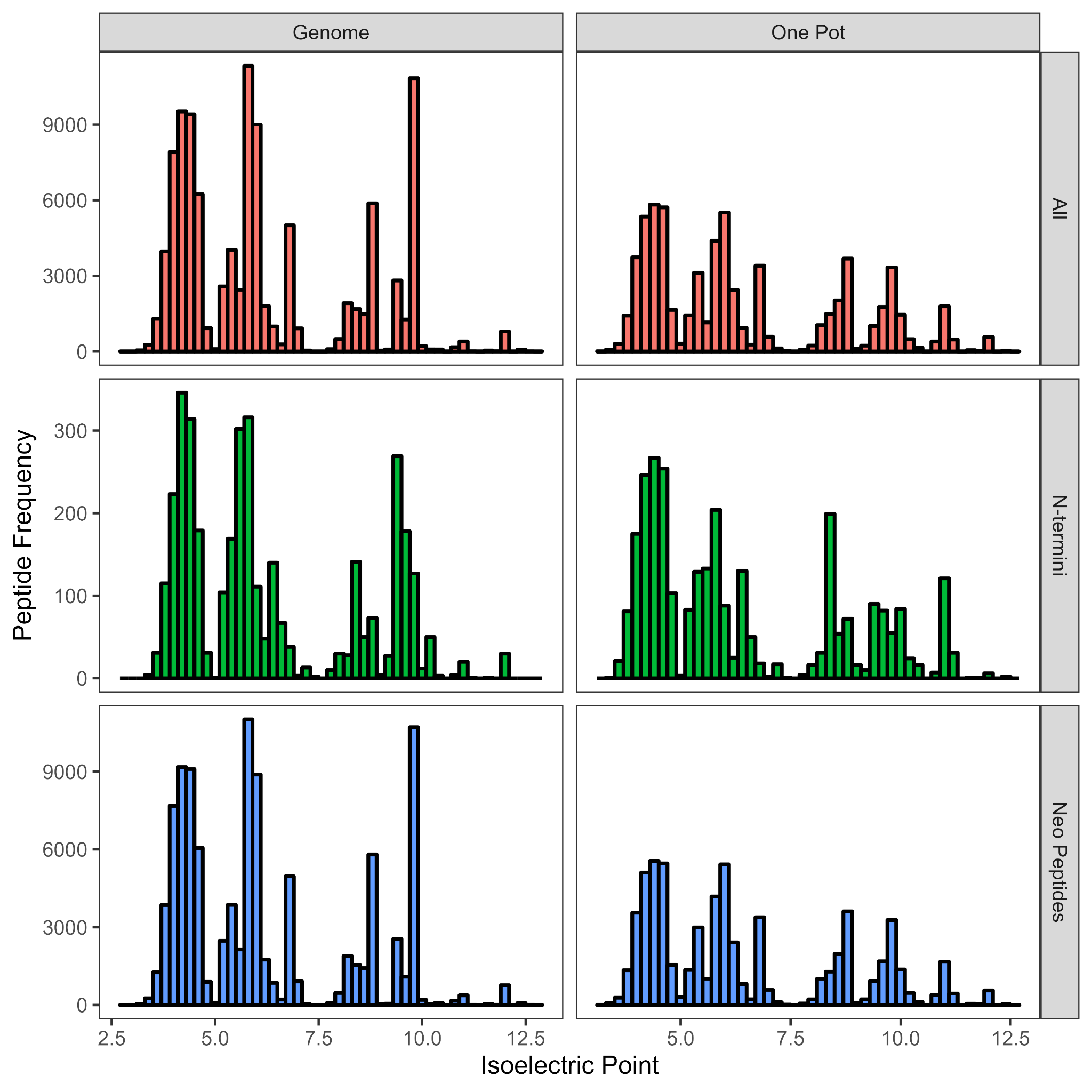

**Supplemental Figure 4.** Histogram of isoelectric points comparing the one-pot method to tryptic peptides generated by the canonical genome. Top panels show all peptides, middle panels show N-terminal peptides, and bottom panels show neopeptides. Left panels are peptides generated from an *in silico* digest of *M. marinum* proteins, whereas right panels are peptides generated from our OnePotNTA technique.

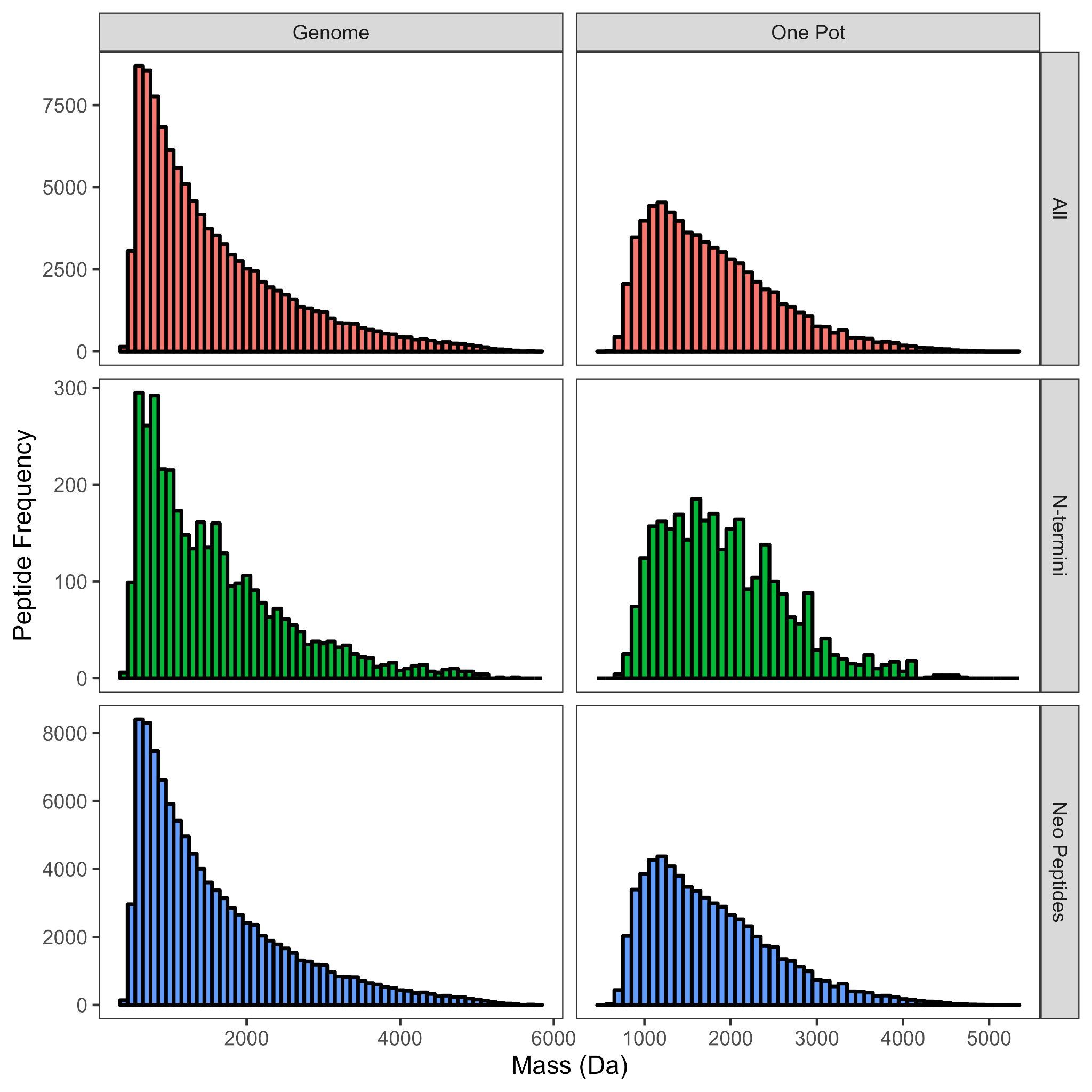

**Supplemental Figure 5.** Histogram of peptide masses between OnePotNTA and tryptic peptides generated by the canonical genome. Top panels show all peptides, middle panels show N-terminal peptides, and bottom panels show neopeptides. Left panels are peptides generated from an *in silico* digest of *M. marinum* proteins, whereas right panels are peptides generated from our OnePotNTA technique.

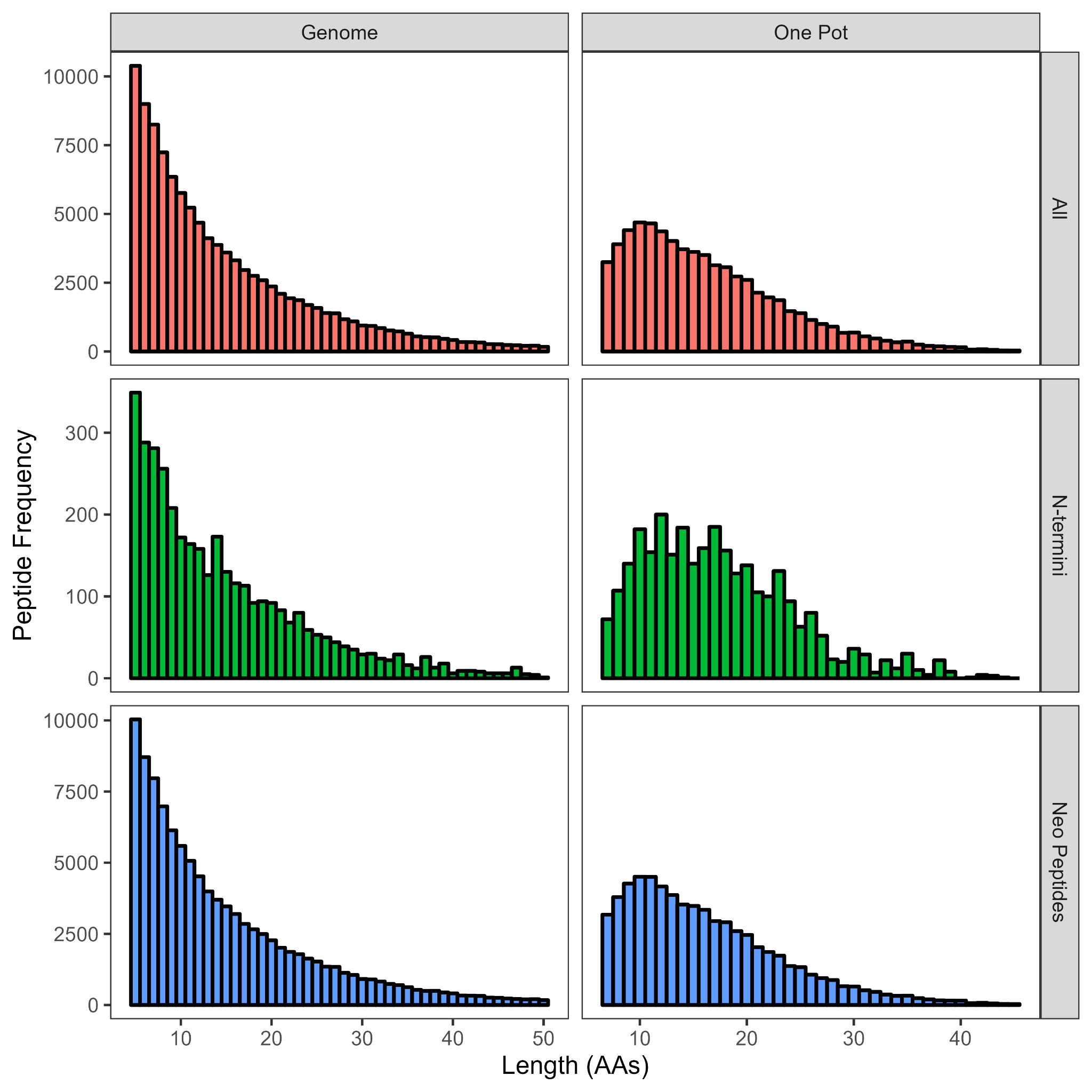

**Supplemental Figure 6.** Histogram of peptide lengths between OnePotNTA and tryptic peptides generated by the canonical genome. Top panels show all peptides, middle panels show N-terminal peptides, and bottom panels show neopeptides. Left panels are peptides generated from an *in silico* digest of *M. marinum* proteins, whereas right panels are peptides generated from our OnePotNTA technique.

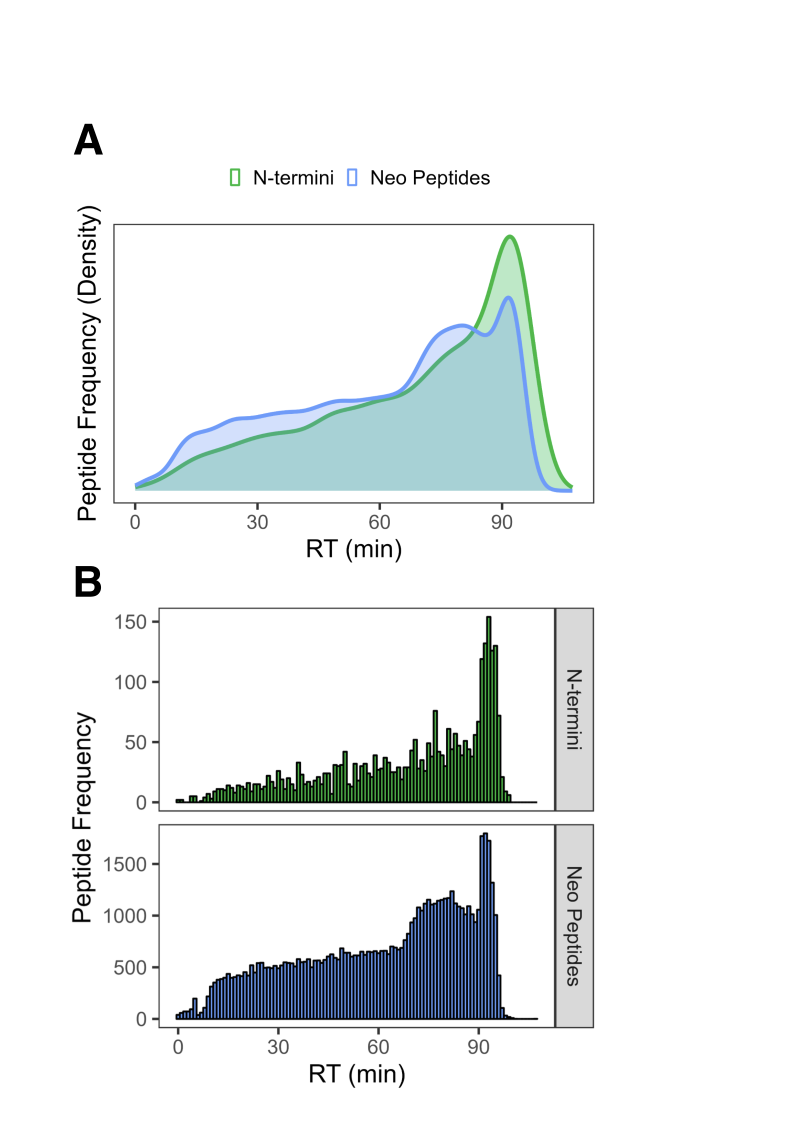

**Supplemental Figure 7**. Distribution of peptide retention times with OnePotNTA. Peptides were separated between N-termini (green) and neopeptides (blue). Panel A shows a density plot using relative intensity (density). Panel B shows a histogram with absolute intensity.

| **MMAR #** | **Gene Name** | **N-terminus** | **MMAR #** | **Gene Name** | **N-terminus** |
| --- | --- | --- | --- | --- | --- |
| MMAR_5363 | accD4_1 | **M**TDTAPGP | MMAR_4683 | Pdc | **M**TLLENDA |
| MMAR_3360 | accD6 | MTIMAPEA | MMAR_2088 | thrS | MSVPAQPA |
| MMAR_4087 | atpD | MTVTAEKT | MMAR_5477 | trxB2 | MTTDSSAD |
| MMAR_4643 | citA | MTVVPEDF | -- | MMAR_1428 | MTESPESP |
| MMAR_1651 | ctaD | **M**TAEAPPL | -- | MMAR_1755 | MSIATNAP |
| MMAR_2677 | eccD5 | MTAVADAP | -- | MMAR_1973 | MTELTQTS |
| MMAR_5450 | esxA | MTEQQWNF | -- | MMAR_2704 | **M**TDMDAAR |
| MMAR_2675 | esxN | MTINYQFG | -- | MMAR_3349 | MTAIDGGT |
| MMAR_4454 | esxN_1 | MTINYQFG | -- | MMAR_3388 | **M**TAGGNPE |
| MMAR_5119 | esxN_2 | MTINYQFG | -- | MMAR_3586 | **M**SVDVTDV |
| MMAR_3653 | esxN_3 | MTINYQFG | -- | MMAR_4583 | MTATSDDS |
| MMAR_3659 | esxN_4 | MTINYQFG | -- | MMAR_4871 | **M**TSDASQD |
| MMAR_3662 | esxN_5 | MTINYQFG | -- | MMAR_5182 | MSDSPSVF |
| MMAR_2289 | fabG1 | **M**TDTATEQ | -- | MMAR_5212 | MTLSLSNH |
| MMAR_0409 | fadD5 | **M**TAELASH | -- | MMAR_5350 | **M**SEEVQSP |
| MMAR_4812 | idi2 | MTNDPAEI | -- | MMAR_5484 | MTEADITE |
| MMAR_2374 | ilvA | **M**SAELSQT |  |  |  |
| MMAR_0057 | leuS | **M**TESPTTT |  |  |  |

**Supplemental Table 1**. Table of genes identified with loss or attenuation of N-terminal acetylation in ∆*emp1* and restoration in complement. **Bold M** = non-canonical initiation (e.g. V, L)

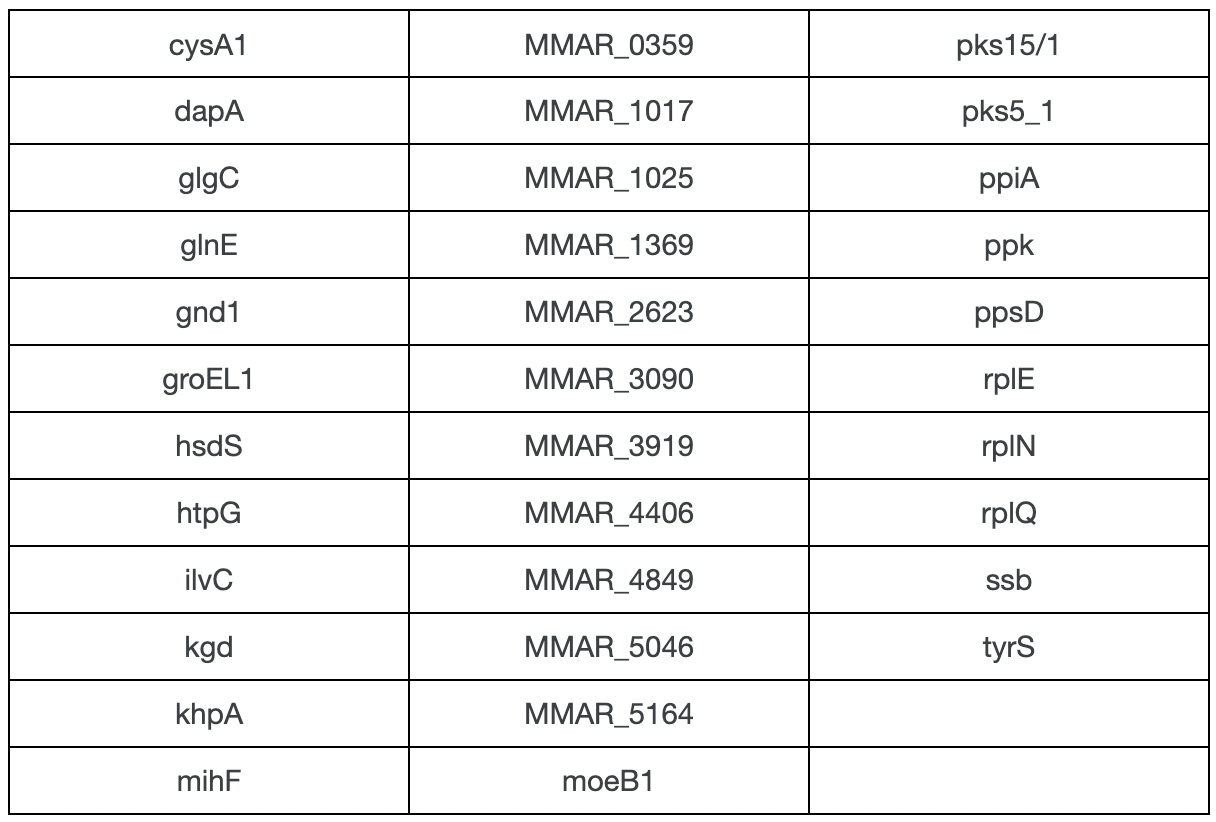

**Supplemental Table 2**. Table of proteins identified with putative noncanonical N-termini. Proteins shown have N-terminal peptides identified with light N-terminal acetylation identified in at least two of three technical replicates in wild type. N-termini begin at position 3 or later in the protein.

**OnePotN𝛼TA Protocol**

Protocol for the labeling and digestion of proteins to quantify protein N-terminal acetylation.

Jump to	:	Stocks	Procedure	Appendix

This protocol is for the labeling and digestion of protein mixtures for bottom-up proteomic analysis of protein N-terminal acetylation. Samples of 100 µg protein are precipitated in acetone and resuspended in buffer and SDS to resolubilize. Precipitation normalizes the variable starting volumes from cell lysates and supernatants.

- Labeling and digestion take place in a commercial S-Trap Mini, which supports up to 300 µg protein per sample. Adaptation to Midi or higher volume devices will likely work. Alternative formats like Miniprep Assisted Proteomics (MAP) will also work.^1^
- Acetone precipitation is performed to normalize concentrations of protein prior to labeling and digestion. Samples with variable volumes (of equal µg of protein) have >7x cold acetone added for up to 1,300 µL total volume. Because of that, limit starting volumes to under 185 µL. Samples with higher starting volume need to be vacuum concentrated (SpeedVac) ahead of time.

**Stocks**

For details on “x##, target ##”, see Magic Numbers

**100 mM TEAB**

*100 mM triethylammonium bicarbonate in water*

- Make ahead in 20 mL scintillation vial.
- 2 mL 1 M TEAB, 18 mL MS water.
- Store @ +4˚C.

**1 M TEAB**

*1 M triethylammonium bicarbonate in water*

- Aliquot taken from commercially available 1 M reagent, to reduce contamination.
- Aliquot in 20mL scintillation vial.
- Store @ +4˚C.

**50 mM ABC**

*50 mM ammonium bicarbonate in water*

- Make ahead in 20 mL scintillation vial.
- 2 mL 500 mM ABC, 18 mL MS water.

**90/10 MeOH/ABC**

*50 mM ammonium bicarbonate in 90% methanol, 10% water*

- Make ahead in 20 mL scintillation vial.
- 2 mL 500 mM ABC, 18 mL MS MeOH.

**90/10 MeOH/TEAB**

*100 mM triethylammonium bicarbonate in 90% methanol, 10% water*

- Make ahead in 20 mL scintillation vial.
- 2 mL 1M TEAB, 18 mL MS MeOH.
- Store @ +4˚C.

**20% SDS**

*20% sodium dodecyl sulfate in water*

- Make ahead in 20 mL scintillation vial.
  1. Weigh 4 g SDS powder.
  2. Add 10 mL MS water.
  3. Cap, place on shaker until fully dissolved (hours).
  4. Add MS water until volume (meniscus) matches another vial with 20 mL of water.
- Store @ RT.

**12% H_3_PO_4_**

*12% phosphoric acid in water*

- Make ahead in 20 mL scintillation vial.
- 17.18 mL water, 2.82 mL 85% phosphoric acid.
- Store @ RT.

**1:1 MeOH/CHCl_3_**

*50% chloroform in methanol*

- Make ahead in 20 mL **amber** scintillation vial.
- 10 mL MeOH, 10 mL CHCl_3._

**90/10 ACN/MeOH**

*10% methanol in acetonitrile*

- Make ahead in 20 mL **amber** scintillation vial.
- 2 mL MeOH, 18 mL MeOH_._

**Q_FA_**

*0.1% formic acid in water*

- Make ahead in 20 mL scintillation vial.
- 20 mL MS water; add 20 µL formic acid (in fume hood).
- Store @ RT.
- Formic acid in +4˚C.

**E_FA_**

*0.1% formic acid, 50% acetonitrile in water*

- Make ahead in 20 mL scintillation vial.
- 10 mL MS water, 10 mL MS acetonitrile; add 20 µL formic acid (in fume hood).
- Store @ RT.

**100 mM TCEP**

*100 mM Tris(2-carboxyethyl)phosphine in 100 mM triethylammonium bicarbonate*

- Make fresh daily in 1.5 mL Epi tube.
- x35, target 14.2, dilute in 100 mM TEAB.

**100mM IAM**

*100mM iodoacetamide in 100mM triethylammonium bicarbonate*

- Make fresh daily in 1.5 mL Epi tube.
- x54, target 9.2, dilute in 100 mM TEAB.

**Resuspended Trypsin^2^**

- Requires 5 µg/5 µL aliquots of trypsin (Promega Gold) in -20˚C already.
- Make right before using.
- +395 µL 50 mM ABC to 5 µg/5 µL trypsin aliquot.
- Makes for 1 µg/80 µL trypsin.

**Resuspended NHS Esters (NHSe)**

- 50 mg aliquots (see Procedure below for synthesis).
- +518 µL 90/10 ACN/MeOH.
- Make right before using.

**Procedure**

- When called to **Add**, vortex and spin down the tube after adding unless stated otherwise.
- When called to **Wash**, aliquot the stated volume into the spin filter inside the tube. Centrifuge for either 1,200g for 30 seconds (remember to balance) or use a countertop “minispin” centrifuge until liquid is visibly passed through. Collection liquid at the bottom needs to be periodically removed (flick into waste).
- When called to **Elute**, proceed like **Wash** but keep all eluate, as that is the final sample.
- Set your heating block to 60˚C in advance to avoid waiting for temperature!

**NHS Ester Synthesis**

*In a rotovap-compatible round-bottom flask*

1. Weigh 1.77 g N-hydroxysuccinimide into flask.
2. Add 5.0 g acetic anhydride-d_6_ from new ampule. Use glass Pasteur pipette.
3. Add small magnetic stir bar. Cover flask neck with foil (loosely).
4. Put flask on cork ring on stir plate. Stir overnight (12-16 hrs).
5. Remove foil cover, stir for additional 12 hours.
6. Pasteur pipette 10 mL hexanes into flask.
7. Attach flask to rotovap, gently evaporate to dryness (no heat).
   *Repeat steps 6 and 7 two more times*
8. Add 7.0 mL anhydrous acetonitrile to flask. Stir to completely dissolve product.
9. Micropipette 141.9 µL (50 mg) aliquots into Epi tubes.
10. SpeedVac aliquots until fully dry
11. Store in -20˚C.

**Precipitation**

*In a 1.5mL tube*

12. Aliquot 100 µg lysate
13. Add cold acetone to total 1,300 µL of volume.

*2 hours @ -20˚C (up to overnight)*

14. Spin 10 mins @ 12,000 g, +4˚C
15. Decant tube, retaining pellet^3^
16. SpeedVac until dry, no heat
17. Add 10 µL 100mM TEAB

**Digest**

*In a 1.5mL tube (same one from precipitation)*

18. Add 5 µL 1M TEAB
19. Add 25 µL 20% SDS
20. Add 5 µL 100mM TCEP

*Vortex to fully dissolve* ^4^

*10 mins @ 60˚C*

*Cool tube to RT* ^5^

21. Add 5 µL 100mM IAM

*30 mins @ dark*

22. Add 5 µL 12% H_3_PO_4_
23. Add 350 µL 90/10 MeOH/TEAB^6^

*Load S-Trap Mini* ^7^

24. Wash 150 µL 90/10 MeOH/TEAB
25. Wash 150 µL 1:1 MeOH/CHCl_3_
26. Add 75 µL NHSe (DO NOT SPIN)
    *30 mins @ RT
    Spin down liquid*
27. Add 75 µL NHSe (DO NOT SPIN)
    *30 mins @ RT
    Spin down liquid*
28. Wash 150 µL 90/10 MeOH/ABC
29. Wash 150 µL 90/10 MeOH/ABC

*New collection tube* ^8^

30. Add 160 µL resuspended trypsin
    (do not fully spin down).

*Mini-spin and Re-top* ^9^

*4 hrs – overnight @ 37˚C*

*Spin down*

31. Elute 80µL Q_FA_
32. Elute 80µL Q_FA_
33. Elute 80µL E_FA_

*20 mins SpeedVac* ^10^

-Samples are now ready for desalting.

**Appendix**

**Explanation for “Magic Numbers”**

Our lab often creates fresh stocks in 1.5mL microcentrifuge (“Epi”) tubes with target concentrations. To simplify the weighing-calculation-dilution process, use these “magic numbers” to simplify the process:

1. “100 mM TCEP” has the numbers “x35, target 14.2”
2. At the balance, weigh your sample as close to 14.2 mg as possible with one scoop. Quick is key, *not* precision.
3. Whatever number you weigh (in mg), multiply it by 35. The resulting number is how much to dilute (in µL) to get the concentration (100 mM).
   a. The “target” value is exactly how much weight is needed to get 500 µL of stock.
   b. 3x the “target” value is the maximum do-not-exceed, as your tube can only fit 1.5 mL

An example formula is provided below for TCEP (x35, target 14.2).

$$\left( \boldsymbol{TCEP wt}, approx 14.2 mg \right)*\left( \boldsymbol{35}\left( \frac{\mu L}{mg} \right) \right)=Suspension Volume for 100 mM$$

**Endnotes**

(1) Mousseau, C. B.; Pierre, C. A.; Hu, D. D.; Champion, M. M. Miniprep Assisted Proteomics (MAP) for Rapid Proteomics Sample Preparation. Analytical Methods, 2023, 15, 916–924. https://doi.org/10.1039/d2ay01549h.
(2) Any trypsin concentration from 1:25 to 1:100 (w/w, enzyme : protein) works for digestion. This protocol sticks to 1:50. Maintain 80 µL volume if digesting with S-Trap Micros (enough to cover the narrowing of the filter), and 160 µL volume if digesting with S-Trap Minis. This is to prevent full evaporation of digestion solution during incubation.

    To make trypsin aliquots, resuspend one vial of Promega Trypsin Gold (V5280) in 100 µL of 0.2% acetic acid (1.0 mL MS water, 2 µL glacial acetic). Hand-flick to dissolve (do not vortex). Portion 5 µL aliquots in 0.5 mL Epi tubes.
(3) Load in tubes centrifuge with cap hinges facing outward. After centrifugation, pellets will retain in-line with the hinge. Decant by tipping the tubes with the hinge facing upwards. This minimizes pellet loss in the decant. Visually inspect that your protein pellet doesn’t delaminate and slip out of the tube when you decant.
(4) This step takes the longest hands-on. Repeatedly vortex, spin down, and visually inspect the bottom of tube to make sure pellet spot is fully dissolved and no longer visible. Apply brief 60˚C heat if necessary. After full dissolution, then proceed with the 10 mins of heat.
(5) Use benchtop “minispin” centrifuge to cool tubes quickly.
(6) This step is known as “flocculation,” the formation of a fine particulate suspension of proteins. At >100 µg protein, solution should look cloudy with tiny flecks of white.
(7) *Load* On S-Trap Micros, this will take two additions of 200 µL, with spin-down in between. Micros don’t hold enough volume for the entire suspension to go in at once.
(8) *New collection tube* Throw away old waste collection tube and place S-Trap in a new 2.0 mL tube, preferably with lid that can close with the filter attached.
(9) *Mini-spin and Re-top* Use the benchtop centrifuge to pulse the spin filters very briefly–1.0 sec. There should be liquid above the filter and a tiny drop of liquid in the collection tube. Use a p200 to “re-top” the bottom liquid back into the filter atop. This step is to ensure trypsin is already permeated throughout the filter before the incubation starts.
(10) After digestion and elution, a final 20 minutes of vacuum concentration is done to reduce/eliminate the acetonitrile used in elution. This prepares your sample (fully aqueous) for desalting via SPE column.
